# Dietary selenium deficiency drives sex-specific circadian disturbance through redox imbalance and causes early systolic dysfunction in mice

**DOI:** 10.64898/2026.09.28.754959

**Authors:** Xuanxuan Guo, Ferdy Lambregtse, Lotte Geerlings, Jumo Zhu, Sietske N. Zijlstra, Anna M. Feringa, Elisabeth M Schouten, Herman H.W. Silljé, Lutz Schomburg, Peter van der Meer, Nils Bomer

## Abstract

**Background:** Selenium is a vital trace element involved in antioxidant defence and cardiovascular health. Although selenium deficiency is implicated in cardiomyopathies, its early cardiac effects and underlying mechanisms remain poorly defined.

**Methods:** C57BL6/Njr mice were fed either a selenium-deficient or control diet for 12 weeks. Systemic selenium status, cardiac function by echocardiography, left ventricular (LV) transcriptomic profiles, redox balance, and circadian pathway markers were assessed, including sex-specific analyses.

**Results:** Selenium deficiency reduced plasma selenium levels without inducing overt cardiac hypertrophy or fibrosis. Echocardiography showed preserved ejection fraction and fractional shortening but reduced global longitudinal strain, indicating early systolic dysfunction. Cardiac stress markers were increased predominantly in male mice. Left ventricular RNA sequencing revealed enrichment of pathways related to cardiac remodelling, redox regulation, mitochondrial function, and circadian rhythm. Additional protein and metabolic analyses supported sex-specific redox–circadian alterations, with males showing a more pronounced stress-response profile.

**Conclusions:** Dietary selenium deficiency induces early myocardial dysfunction and molecular remodelling before overt cardiac failure. These changes are associated with redox and circadian pathway disruption and show sex-specific features, suggesting that selenium contributes to cardiac homeostasis through sex-dependent redox–circadian regulation.

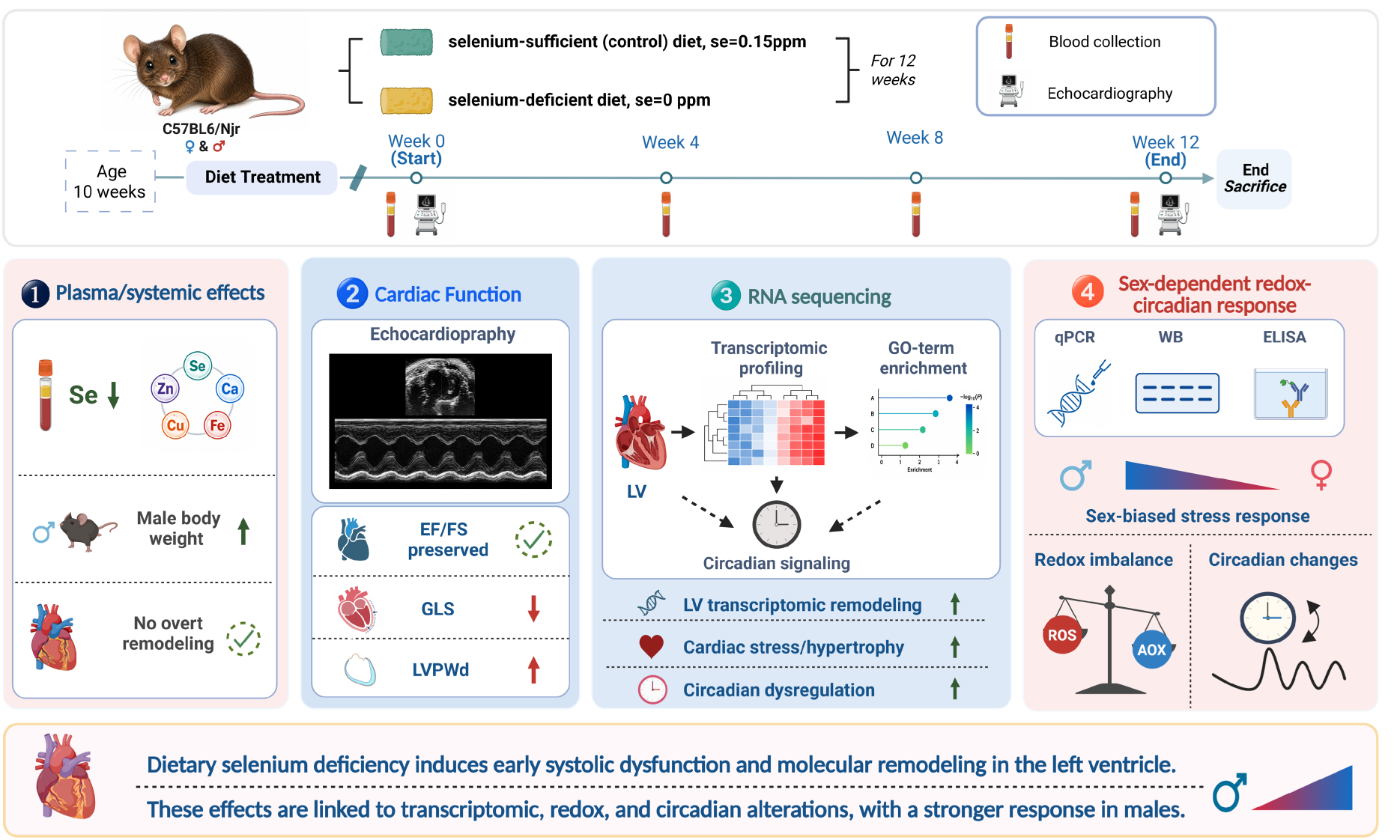

**TRANSLATIONAL PERSPECTIVES:** *What Is New?:*

- Dietary selenium deficiency induces early myocardial dysfunction before overt heart failure develops, as detected by impaired global longitudinal strain despite preserved ejection fraction.
- Selenium deficiency promotes cardiac molecular remodelling characterized by activation of hypertrophic signalling, mitochondrial stress, and disruption of interconnected redox and circadian pathways.
- The cardiac response to selenium deficiency is sexually dimorphic, with male mice exhibiting greater oxidative stress, circadian dysregulation, and activation of cardiac stress pathways than females.

*What Are the Clinical Implications?:* Selenium deficiency induces early myocardial dysfunction before overt cardiac failure, accompanied by activation of hypertrophic gene programmes and disruption of redox and circadian homeostasis. These effects are more pronounced in male mice, suggesting sex-specific mechanisms of susceptibility. The findings identify a redox–circadian axis as a potential mechanistic link between selenium deficiency and cardiac remodelling and support evaluation of selenium status as a modifiable cardiovascular risk factor. Early nutritional intervention and therapies targeting redox–circadian signalling may offer new opportunities to prevent progression to clinically overt heart failure.

## 1. INTRODUCTION

Heart failure (HF) is a complex, life-threatening clinical syndrome in which the heart is unable to pump blood sufficiently to meet the metabolic demands of the body[1]. It primarily affects the older population, and with increasing life expectancy and improved survival from cardiovascular diseases, the global prevalence of HF is expected to rise substantially in the coming decades[2]. This trend presents a growing clinical and socioeconomic burden. HF has a multifactorial aetiology, including myocardial infarction, hypertension, valvular heart disease, and cardiotoxic chemotherapy[3]. However, in many cases, no specific cause can be identified.

Emerging evidence highlights the role of nutritional factors in HF pathophysiology, particularly the imbalance of circulating minerals and trace elements such as iron, iodine, zinc, and selenium[4–7]. Up to 50% of patients with HF exhibit some form of malnutrition, including micronutrient deficiencies, which are increasingly recognized as contributors to disease progression[8, 9]. Selenium, an essential micronutrient with critical antioxidant and immunomodulatory functions, has been implicated in cardiomyopathy. A well-documented example is Keshan disease, a rare but fatal form of dilated cardiomyopathy (DCM) found in selenium-deficient regions of China [4, 10]. This condition shares clinical features with idiopathic DCM and was virtually eradicated through government-led selenium supplementation programmes, underscoring selenium’s role in cardiovascular health [11].

Despite this, the precise mechanisms by which selenium deficiency contributes to HF remain poorly understood. Proposed pathways include oxidative stress, impaired selenoprotein function, and modulation of host susceptibility to viral infections such as coxsackievirus B[12–14]. More recent data from the BIOSTAT-CHF cohort, a large European study of patients with worsening HF, showed that approximately 25% of patients had selenium deficiency (serum Se <70 µg/L)[15]. These patients, particularly older women, exhibited worse quality of life, reduced exercise capacity, and poorer clinical outcomes, suggesting a strong association between low selenium levels and HF severity. Furthermore, in the Dutch general population cohort PREVEND (N=∼6000) it was shown that having sufficient selenium levels was associated with reduced risk for new-onset HF and similarly for new-onset AF[16].

Given these findings, we hypothesize that selenium deficiency contributes directly to the development and progression of HF. In this study, we aim to investigate the mechanistic relationship between selenium deficiency and HF pathophysiology using a diet-induced selenium-deficient mouse model. By examining the effects on myocardial function and delineating the involved molecular pathways, we seek to provide new insights into the role of selenium in HF and explore potential therapeutic strategies targeting micronutrient deficiencies.

## 2. MATERIALS AND METHODS

### 2.1 Animal experiments

All animal studies were approved by the Central Committee of Animal experiments (De Centrale Commissie Dierproeven, CCD, licence number AVD10500202115446) and the Animal Welfare Body of the University of Groningen (permit number 2115446-02-003). And this study followed the guidelines set out in the Directive 2010/63/EU of the European Parliament for using animals in research which were reported in line with the research: The ARRIVE guidelines 2.0: updated guidelines for reporting animal research [17]. C57BL6/Njr mice were provided by Janvier Labs and housed in Centrale Dienst Proefdieren (CDP) facility of the UMCG under a 12 h light/12 h dark cycle with *ad libitum* access to chow and water. Animals were allowed to acclimatise to the animal facility for 1 week before initiation of the dietary intervention.

#### 2.1.1 Selenium controlled diet treatment, body weight measurement and plasma isolation

To investigate the intrinsic and systemic effects of dietary selenium status on the heart, selenium-controlled dietary interventions were initiated when the mice were 10 weeks old. C57BL6/Njr mice were allocated to either a selenium-deficient diet group (0.013 ppm Se; Research Diets, Inc., lot A23050903) or a selenium-sufficient control diet group (0.15 ppm Se; Research Diets, Inc., lot A23050905), with 10 male and 10 female mice included in each dietary group. Group size was informed by an a priori power calculation performed using G*Power version 3.1.9.4. Before dietary allocation, all mice underwent baseline echocardiography and were confirmed to have normal baseline ejection fraction. Mice were first stratified by sex and subsequently allocated to the two dietary groups while balancing baseline body weight between groups. Body weight was measured on the first day of dietary intervention and weekly thereafter. To monitor changes in plasma trace element concentrations, blood samples were collected at weeks 0, 4, 8, and 12 by orbital blood sampling under anaesthesia, with no more than 10% of the estimated circulating blood volume collected at each time point.

Anaesthesia was administered as described below. The individual mouse was considered the experimental unit. The order of treatments, sample collection, and measurements was alternated between groups to minimise potential order effects.

Blood was collected in EDTA tubes and centrifuged at 1500 × g for 10–15 min at room temperature. The supernatant (EDTA plasma) was transferred to a new 1.5 mL tube, initially stored in liquid nitrogen, and subsequently transferred to −80 °C freezers for long-term storage.

#### 2.1.2 Anaesthesia and analgesia

Mice were anaesthetised with 2.0–2.5% isoflurane in oxygen, administered continuously by inhalation via a mouthpiece during echocardiography, orbital blood sampling, and terminal procedures.

Anaesthesia was maintained throughout each procedure. No analgesic agents were administered.

#### 2.1.3 Euthanasia and terminal tissue collection

After 12 weeks of dietary treatment, euthanasia and terminal tissue collection were performed under continuous deep isoflurane anaesthesia. Anaesthesia was maintained with 2.0–2.5% isoflurane in oxygen throughout the entire terminal procedure. Adequate depth of anaesthesia was confirmed by the absence of a pedal withdrawal response to toe pinch. The abdominal cavity was opened, and terminal blood was collected via the abdominal aorta. Fifty microlitres of whole blood were transferred to a separate 1.5 mL tube and snap-frozen in liquid nitrogen. The remaining blood was collected in EDTA tubes, centrifuged, and the resulting EDTA plasma was stored in liquid nitrogen.

Subsequently, the thoracic cavity was opened and the circulatory system was flushed with saline via cardiac apical puncture. Euthanasia was completed by excision of the heart under continuous deep anaesthesia, after which isoflurane administration was discontinued.The excised heart was rinsed in ice-cold 1 M potassium chloride (KCl). The right ventricle (RV) and atria were dissected from the left ventricle (LV), weighed, and stored separately in liquid nitrogen. A mid-ventricular transverse section of the LV was fixed in 4% formalin before paraffin embedding, as described previously [18, 19], while the remaining LV tissue was snap-frozen for molecular analyses. The liver, spleen, and kidneys were also excised and weighed. Portions of these tissues were fixed in 4% formalin and subsequently processed for paraffin embedding, while the remaining tissue was snap-frozen for molecular analyses. Tibiae were also collected at the terminal time point.

### 2.2 Plasma Trace element concentration measurements

Aliquots of 50 µL EDTA plasma samples were sent to Charité Universitätsmedizin Berlin for quantitative analysis. To assess the impact of a selenium-deficient diet on selenium levels in mice, total reflection X-ray fluorescence (TXRF; S4 T-STAR, Bruker Nano GmbH, Berlin, Germany) was used to determine the concentration of Ca, Cu, Fe, Se and Zn. Samples were diluted with a Gallium standard solution for internal control, applied to polished glass plates and analysed by TXRF as described. [20, 21]

### 2.3 Echocardiography

Echocardiography (Echo) was performed by a blinded observer on week 0 (baseline record) and week 12 (after 12-weeks selenium controlled dietary, 2-3 days before sacrificing) to assess cardiac structure and function using the Vevo 3100 system (FuJiFLIM VisualSonics, Toronto, Canada), equipped with 40 MHz MXX550D linear array transducer, as previously described[22]. Mice were anaesthetized with 2–2.5% isoflurane in oxygen during the procedure. Body temperature was maintained at 37 °C. The Vevo LAB software version 5.5.0 (FuJiFLIM VisualSonics) was used to measure echocardiographic parameters.

### 2.4 Masson trichrome staining

All tissues harvested and fixed in 4% formalin in 2.1.2 were processed and prepared according to previously described protocols [18, 19, 23]. Paraffin-embedded tissues were cut into 4 μm sections for histological analyses. Stained sections were automatically scanned using a Nanozoomer 2.0-HT digital slide scanner (Hamamatsu, Japan). The percentage fibrosis per section was determined using Aperio’s ImageScope software and was quantified as percentage of the entire section at 40× magnification.

### 2.5 RNA isolation and quantitative real time polymerase chain reaction (qRT-PCR)

Total RNA was isolated from frozen LV tissue using TRI reagent (Sigma-Aldrich, St Louis, MO) and cDNA was made using the QuantiTect® Reverse Transcription Kit (Qiagen, Germany) according to the manufacturer’s protocol. qPCR was performed using a Bio-Rad CFX384 Real-Time PCR system (Bio-Rad, CA, USA) using SYBR Green dye. Gene 36B4 was used to correct for the measured mRNA expression as the housekeeping gene. Primer sequences are shown in Supplementary Table 4. The selenium sufficient group was used as the control group.

### 2.6 RNA sequencing

RNA sequencing was outsourced to Lexogen GmbH. A total of 20 samples, derived from two experimental conditions, were prepared and submitted for sequencing. Library preparation was performed using Lexogen’s CORALL library preparation kit. Sequencing was conducted using paired-end reads, and samples included ERCC and SIRV spike-in controls for quality control and normalization purposes. The sequencing data were aligned to the *Mus musculus* reference genome (GRCm38.101) supplemented with ERCC and SIRV annotations. Raw sequencing reads were quality-checked, and adapters were trimmed by Lexogen prior to alignment. Each sample generated between 18 and 64 million read pairs (median: 28.93 million). Reads were aligned to the reference genome using the STAR aligner, which allows for fast and accurate alignment through a pre-built genome index. Uniquely mapped reads ranged from 14.23 to 56.00 million per sample (median: 23.60 million), indicating high-quality alignment performance. Unique Molecular Identifiers (UMIs) were used to collapse PCR duplicates. Position-based and gene-based collapsing approaches were employed to retain only unique reads mapping to the same genomic position or gene, respectively, thereby improving quantification accuracy. Post-alignment quality metrics were assessed using the RSeQC package, which included analyses of read distribution across genomic features (coding sequence, UTRs, introns, and intergenic regions). The read_distribution.py script was utilized to classify reads based on their genomic locations, providing an overview of transcriptomic complexity and potential biases.

RNA-sequencing data were analysed in R (version 4.4.2). Differential gene expression analysis was performed using the DESeq2 package. Raw count data were normalized using the median-of-ratios method, and differential expression between experimental groups was assessed by fitting negative binomial generalized linear models. Statistical significance was determined using Wald tests, and p-values were adjusted for multiple testing using the Benjamini–Hochberg procedure. Genes with an adjusted p-value (padj) < 0.05 and an absolute log2 fold change (|log2FC|) > 1 were considered differentially expressed.

Variance-stabilizing transformation was applied to normalized count data prior to downstream exploratory analyses. Principal component analysis (PCA) was performed to evaluate sample clustering and experimental reproducibility. Data visualization was conducted using the ggplot2 package. Differential expression results were visualized using volcano plots generated with the EnhancedVolcano package. Heatmaps of normalized gene expression were generated using the ComplexHeatmap package following row-wise scaling of expression values.

Functional pathway enrichment analysis of differentially expressed genes was performed using the Kyoto Encyclopedia of Genes and Genomes (KEGG) database. Significantly enriched pathways were visualized using the pathview package, which maps gene expression changes onto curated KEGG pathway diagrams.

### 2.7 Western blotting

Protein level comparisons were performed with Western blot. Frozen left ventricular mice tissue was treated in RIPA buffer (50mM TRIS; 1% I IGEPAL® CA-630; sodium deoxycholate, 0,5% W/V; 0.1% SDS; 0.15 M NaCl; in deionized water) with 40 µl/mL cOmplete™, Mini Protease Inhibitor Cocktail, 10 µl/mL Phosphatase inhibitor cocktail 3, 30 µl/mL NaVanadate, and 5 µl/mL PMSF (200 mM in isopropanol) in the TissueLyser LT system. Protein concentrations were determined with Pierce™ BCA Protein Assay Kits (Thermo Scientific™, cat no. 23225), using Gen5 2.04 on SynergyH1 plate reader. Samples were run in a 12% SDS-PAGE gel 15 min at 70 mVa/ 90 min at 120 mVa. Semi-dry Western blotting onto PVDF membrane proceeded for 75 min at 0.8 mA/cm^2^. Prior to staining blots were blocked for 1 hour in 5% powdered milk. Total protein stain was performed with Revert 700 Total Protein Stain (LI-COR, cat no. 926-11021) and blots were imaged on the Amersham™ ImageQuant™ 800 Western blot imaging system (Cytiva, cat no. 29399481) at 700 nm. Blots were stained overnight with respective primary antibody at 4 °C and respective secondary antibody for 1 hour at RT. Between staining steps, blots were washed 3x 5 min with PBS with 1% Tween. Western Lightning Ultra, Chemiluminescent Substrate (revvity, cat no. NEL111001EA) was used for visualization on Amersham™ ImageQuant™ 800 Western blot imaging system (Cytiva, cat no. 29399481) using the IQ800 Control software for chemiluminescence quantification. Blot images were analysed with Image Studio Lite Ver 5.2 and statistical analysis was performed with GraphPad Prism 10.2.3. ROUT test for outliers analysis was applied. Subsequently, grouped analysis was performed with two-way ANOVA and Tukey’s multiple comparisons test, and for non-grouped unpaired t-test. *p < 0.05 was considered significant.

### 2.8 GSH/GSSG ratio determination

To determine the GSH/GSSG ratio the Invitrogen™ Glutathione Colorimetric Detection Kit (Invitrogen, cat no: EIAGSHC) was used according to the accompanying protocol on frozen powder of mice left ventricular tissue.

### 2.9 NAD+/NADH ratio determination

The NAD+ and NADH concentrations were determined using frozen powder mice left ventricular tissue and the Sigma-Aldrich NAD+/NADH Assay Kit (Sigma-Aldrich, cat no. MAK460-1KT) according to the accompanying protocol.

### 2.10 Statistical analysis

Continuous data are presented as mean ± standard error of the mean (SEM), unless stated otherwise. Individual values are shown where applicable. Statistical analyses were performed using GraphPad Prism (v.10.2.3), except for RNA sequencing analyses, which were performed using DESeq2 in R (v.4.2.0). Comparisons between experimental groups were performed using unpaired two-tailed Student’s t-tests, Mann–Whitney tests, one-way ANOVA, or two-way ANOVA, as appropriate. Post-hoc multiple comparisons were performed where applicable. For analyses involving diet and sex as factors, two-way ANOVA was used to assess main effects and interactions. For RT-qPCR analyses, statistical tests were performed on ΔCt values, whereas fold-change values were calculated relative to the corresponding control group for presentation. Fold changes for Western blot analyses were calculated relative to the corresponding control group, as indicated in the figure legends. Outlier analysis was performed using the ROUT method (Q = 1%) where applicable. For RNA sequencing, adjusted P-values were calculated using the Benjamini–Hochberg false discovery rate method, with Padj < 0.05 considered statistically significant. For downstream interpretation and enrichment analyses, genes were additionally filtered using an absolute log2 fold-change threshold of >0.5. For all other analyses, P < 0.05 was considered statistically significant.

## 3. RESULTS

### 3.1 Selenium deficient diet resulted in decreased plasma selenium concentration, and accelerated body weight gain in male mice

Selenium-deficient diet induced systemic selenium deficiency in mice within 12 weeks, as evidenced by the drastically reduced plasma selenium concentrations compared to the control group (Figure 1B-C). No significant effects were observed on plasma iron and zinc concentrations. Plasma copper levels were slightly decreased in the female selenium-deficient group (Supplemental Table 1). Additionally, male mice under the selenium deficient regime gained significantly more body weight compared to the male controls, but this was not observed between the female groups (Figure 1D). Assessing cardiac weights (LV, RV, atria and whole heart) selenium deficiency did not induce overt cardiac hypertrophy (Figure 1E). Also, staining did not show detectable fibrotic remodelling in the left ventricle or other major organs (Supplemental Tables 2–3).

**Figure 1:**
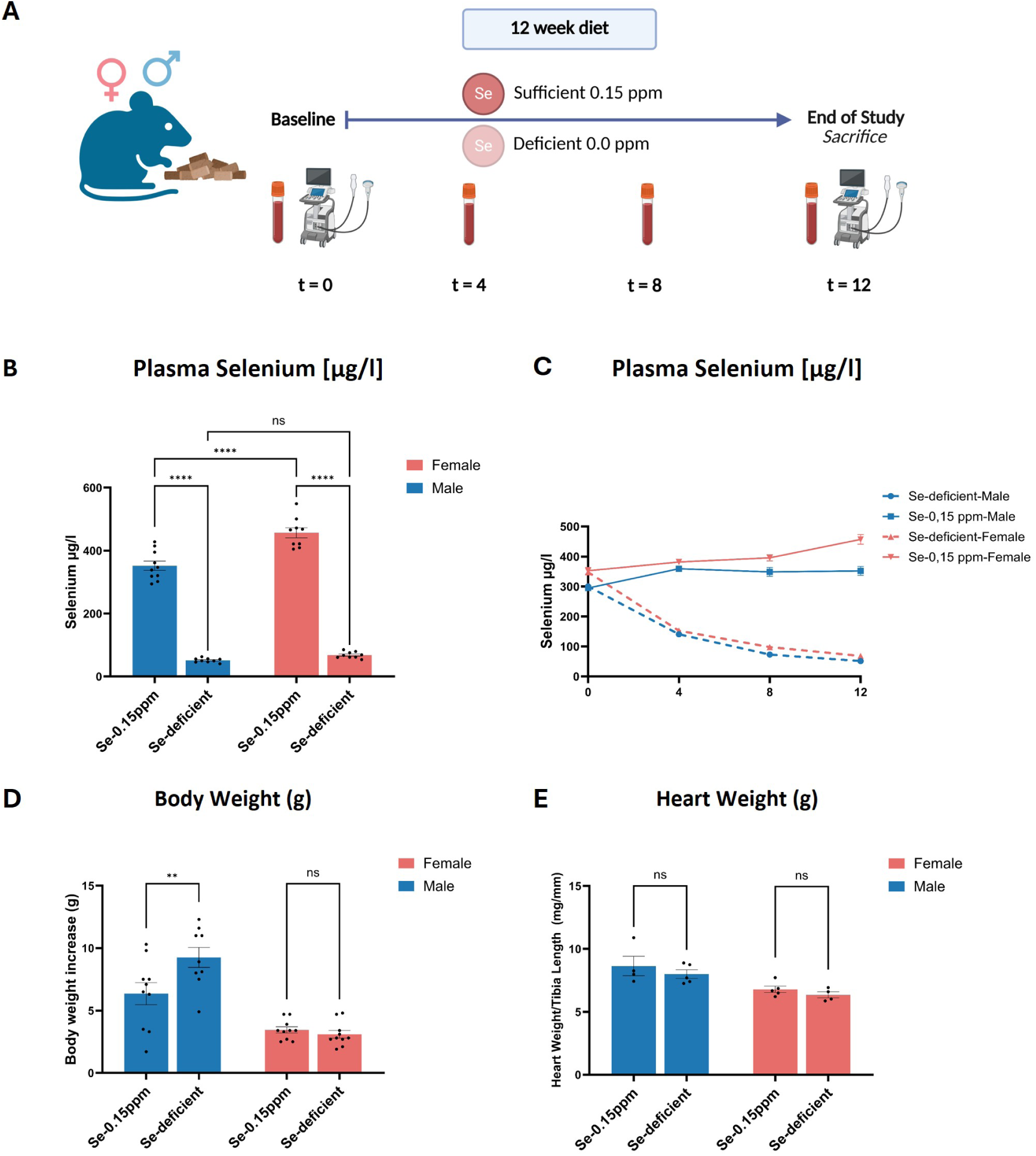
Systemic effects of a selenium-deficient diet in mice. **A)** Schematic representation of the study design, where 10-week-old C57BL6/Njr mice were subjected to either a selenium-sufficient (0.15 ppm) control diet or a selenium-deficient diet for 12 weeks. Echocardiography was performed at baseline and at the end of the study, after which the animals underwent terminal procedures. **B)** Plasma selenium concentration at the end of the study. **C)** Plasma selenium concentration over time during the study period. **D)** Body weight increase (g), separated by sex. **E)** Heart weight corrected for tibia length, separated by sex. Data are presented as individual values with mean ± SEM. For panel B, Se-sufficient males, n = 10; Se-deficient males, n = 9; Se-sufficient females, n = 9; and Se-deficient females, n = 9 independent biological replicates. Statistical significance was assessed using two-way ANOVA followed by Šídák’s multiple-comparisons test. For panel C, for weeks 0, 4, 8, and 12, respectively, the numbers of independent biological replicates were n = 10/9/9/9 for Se-deficient males, n = 10/10/10/10 for Se-sufficient males, n = 10/10/10/9 for Se-deficient females, and n = 10/10/10/9 for Se-sufficient females. For panel D, Se-sufficient males, n = 10; Se-deficient males, n = 9; Se-sufficient females, n = 10; and Se-deficient females, n = 10 independent biological replicates. Selenium-sufficient and selenium-deficient groups were compared separately within each sex using unpaired two-tailed Student’s t-tests. For panel E, Se-sufficient males, n = 4; Se-deficient males, n = 5; Se-sufficient females, n = 5; and Se-deficient females, n = 4 independent biological replicates. Selenium-sufficient and selenium-deficient groups were compared separately within each sex using unpaired two-tailed Student’s t-tests. **P < 0.01, ****P < 0.0001; ns, not significant.

### 3.2 Selenium-deficient diet induced early sex-specific cardiac alterations at functional and molecular levels

No significant differences in ejection fraction (EF) (Control: 54.70 ± 2.17% vs. Se-deficient: 56.30 ± 2.62%, Table 1) or fractional shortening (FS) (Control: 28.23 ± 1.43% vs. Se-deficient: 29.32 ± 1.68%, Table 1) were observed between groups (Figure 2A–B). However, selenium-deficient mice exhibited a significantly impaired global longitudinal strain (GLS) compared to controls (Control: –15.28 ± 0.71% vs. Se-deficient: –12.21 ± 1.00 %, Table 1), indicating early myocardial dysfunction (Figure 2C). In addition, no significant differences were detected in other cardiac parameters, such as heart rate (HR) and left ventricular dimensions. Notably, left ventricular posterior wall thickness in diastole (LVPWd) was significantly increased in the selenium-deficient group (Control: 0.8368 ± 0.02 mm vs. Se-deficient: 0.9429 ± 0.03 mm) (Table 1), suggesting early structural remodelling.

**Figure 2.**
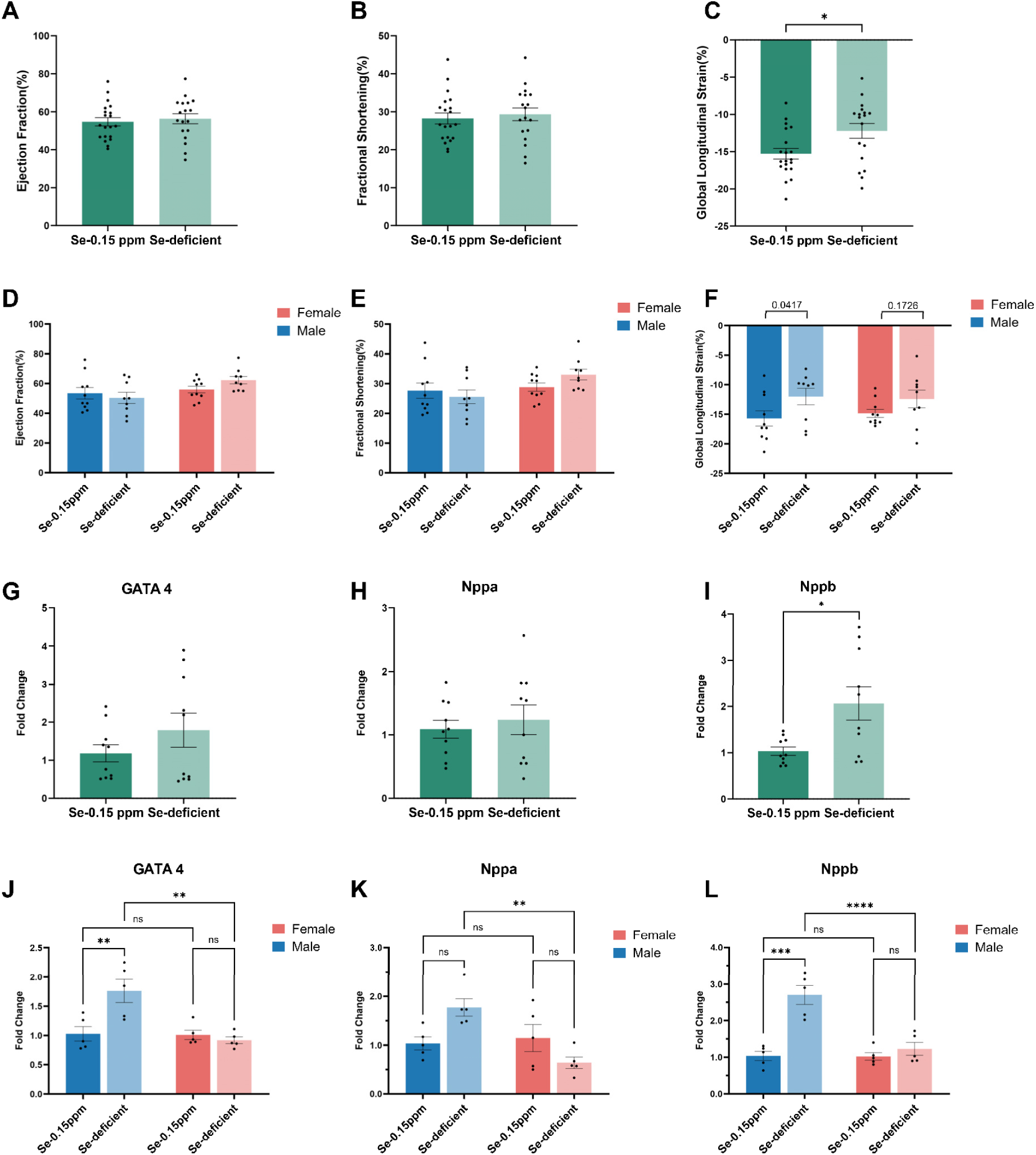
Cardiac functional and molecular assessments in mice. **A)** Left ventricular ejection fraction (EF), **B)** fractional shortening (FS), and **C)** global longitudinal strain (GLS) in mice on the control (Se-0.15 ppm) or selenium-deficient diet at week 12 of the study. **D)** Sex-specific analysis of EF, **E)** FS, and **F)** GLS in male and female mice. RT-qPCR fold changes of **G)** GATA4, **H)** Nppa, and **I)** Nppb. Sex-specific analysis of **J)** GATA4, **K)** Nppa, and **L)** Nppb. Data are presented as individual values with mean ± SEM. For panels A–C, n = 20 selenium-sufficient and n = 18 selenium-deficient mice (independent biological replicates). Selenium-sufficient and selenium-deficient groups were compared using unpaired two-tailed Student’s t-tests. For panels D–F, Se-sufficient males, n = 10; Se-sufficient females, n = 10; Se-deficient males, n = 9; and Se-deficient females, n = 9 independent biological replicates. Data were analysed using two-way ANOVA followed by Šídák’s multiple-comparisons test. In panel F, exact unadjusted P values from uncorrected Fisher’s LSD pairwise comparisons are additionally shown for the within-sex diet comparisons (males, P = 0.0417; females, P = 0.1726); these comparisons did not remain significant after Šídák correction. For panels G–I, n = 10 selenium-sufficient and n = 10 selenium-deficient independent biological replicates. For panels J–L, Se-sufficient males, n = 5; Se-sufficient females, n = 5; Se-deficient males, n = 5; and Se-deficient females, n = 5 independent biological replicates. Each biological sample was measured in technical duplicate by RT-qPCR. Selenium-sufficient and selenium-deficient groups in panels G–I were compared using unpaired two-tailed Student’s t-tests. For panels J–L, statistical analyses were performed on ΔCt values using two-way ANOVA followed by Šídák-adjusted multiple comparisons for the pairwise comparisons shown. *P < 0.05, **P < 0.01, ***P < 0.001, ****P < 0.0001; ns, not significant.

**Figure 3:**
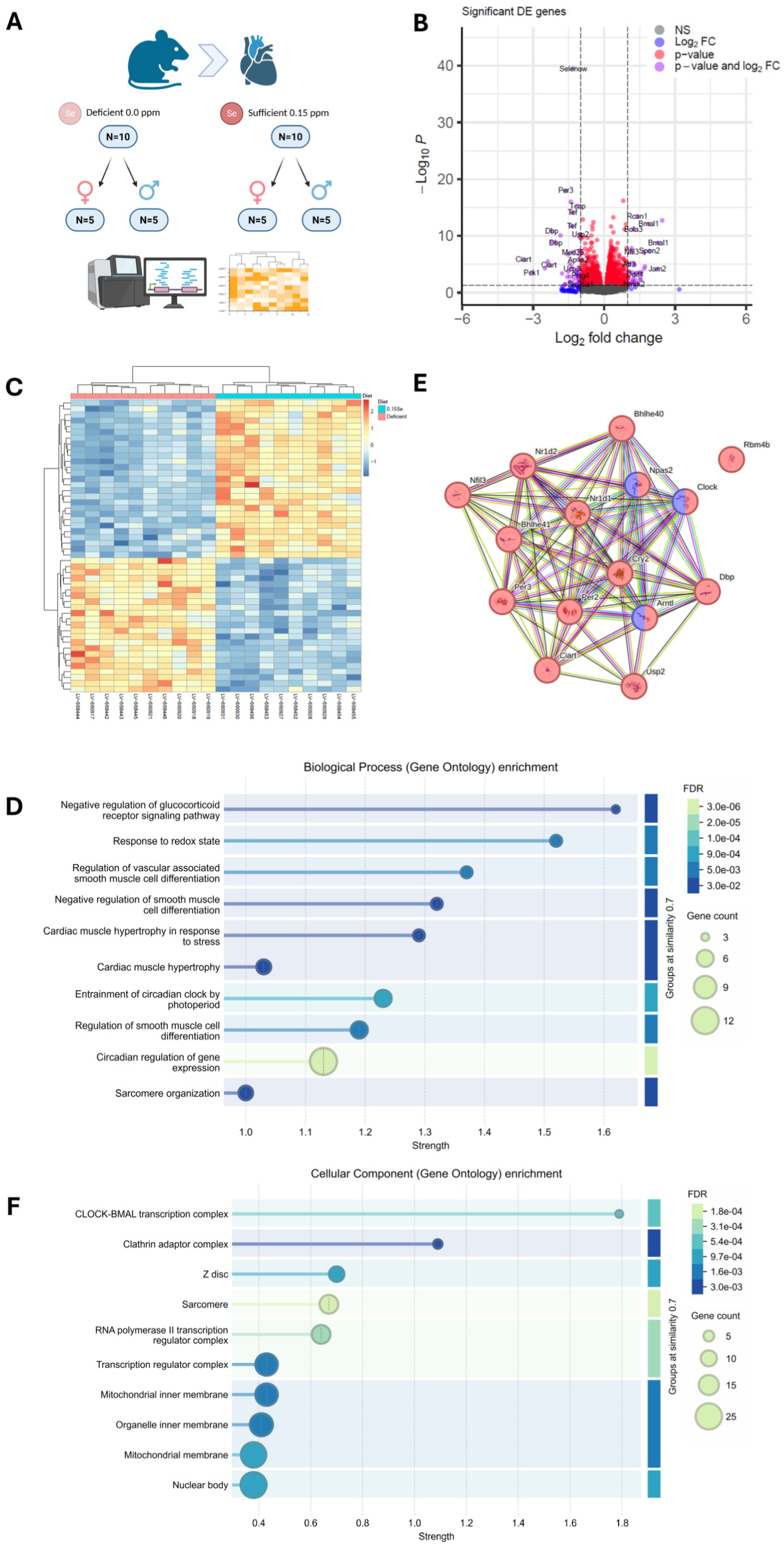
RNA sequencing analyses of left ventricle tissue of selenium-sufficient and selenium-deficient mice. **A)** Schematic overview of the experimental set-up. **B)** A volcano plot derived from the RNA sequencing results. **C)** A heatmap showing the group separation based on the 50 most significant differentially expressed transcripts, without stratifying by sex. **D)** GO-term enrichment for Biological Processes. **E)** Network of identified DEGs involved in circadian rhythm. Nodes coloured red are members of the enriched GO term GO:0007623 related to circadian rhythm, while nodes coloured blue and red are also members of the enriched GO term GO:0051775 related to the response to redox state. **F)** GO-term enrichment for Cellular Components. RNA sequencing was performed using n = 10 mice per dietary group (5 male and 5 female independent biological replicates per group). Differential expression was analysed using DESeq2 with Wald tests and Benjamini–Hochberg correction for multiple testing. GO-term enrichment was performed using clusterProfiler::enrichGO() with hypergeometric over-representation analysis and Benjamini–Hochberg correction; q values < 0.05 were considered significant.

**Table 1.** Echocardiographic parameters. Data are presented as mean ± SEM. Sex-stratified comparisons were analysed using two-way ANOVA followed by Šídák’s multiple-comparisons test. Comparisons between the total selenium-sufficient and selenium-deficient groups were performed using unpaired two-tailed Student’s t-tests. ‡P < 0.05 compared with the total control group. *FS = fractional shortening; EF = ejection fraction; CO = cardiac output; HR = heart rate; LVPWs = left ventricular posterior wall thickness in systole; LVPWd = left ventricular posterior wall thickness in diastole; LVAWs = left ventricular anterior wall thickness in systole; LVAWd = left ventricular anterior wall thickness in diastole; GLS = global longitudinal strain, a marker of myocardial deformation*.

| Echocardiographic parameters | Se-0.15 ppm (control) |  |  | Se-deficient |  |  |
| --- | --- | --- | --- | --- | --- | --- |
|  | Males, n=10 | Females, n=10 | Total, n=20 | Males, n=9 | Females, n=9 | Total, n=18 |
| EF (%) | 53.46±3.81 | 55.94±2.22 | 54.70±2.17 | 50.34±3.75 | 62.27±2.49 | 56.30±2.62 |
| FS (%) | 27.64±2.57 | 28.83±1.41 | 28.23±1.43 | 25.58±2.31 | 33.06±1.79 | 29.32±1.68 |
| CO(mL/min) | 14.92±0.83 | 14.80±0.59 | 14.86±0.50 | 15.62±0.91 | 12.98±0.92 | 14.30±0.70 |
| HR (bpm) | 444.0±13.65 | 427.1±10.43 | 435.6±8.58 | 454.2±14.21 | 410.3±15.16 | 423.3±11.40 |
| LVPWs (mm) | 1.13±0.04 | 1.17±0.04 | 1.15±0.03 | 1.30±0.08 | 1.25±0.08 | 1.28±0.06 |
| LVPWd (mm) | 0.89±0.04 | 0.78±0.02 | 0.8368±0.03 | 0.98±0.05 | 0.91±0.03 | 0.9429±0.03<sup>‡</sup> |
| LVAWs (mm) | 1.37±0.07 | 1.25±0.02 | 1.312±0.04 | 1.36±0.06 | 1.37±0.03 | 1.362±0.03 |
| LVAWd (mm) | 0.92±0.04 | 0.87±0.02 | 0.8914±0.02 | 0.97±0.04 | 0.92±0.04 | 0.9464±0.03 |
| GLS (%) | -15.71±1.29 | -14.86±0.67 | -15.28±0.71 | -12.01±1.40 | -12.42±1.50 | -12.21±1.00<sup>‡</sup> |

When stratified by sex, no significant differences were observed in EF or fractional shortening between selenium-deficient and control groups in either males or females (Figure 2A–B, D–E; Table 1). Notably, a significant main effect of sex was observed, with reduced EF and FS in males and increased EF and FS in females (*P*sex = 0.029 and 0.045, respectively). Nevertheless, no significant sex × diet interaction was observed. Similarly, other conventional echocardiographic parameters remained comparable between sex groups. Sex-stratified analysis of GLS showed a significant effect of diet, but not of sex. However, in unadjusted pairwise comparisons, the diet difference in GLS in males was nominally significant (*P* = 0.0417), while the corresponding difference in females was less pronounced (*P* = 0.1726) (Figure 2F). These within-sex comparisons did not remain significant after correction for multiple comparisons.

To assess whether molecular alterations in the heart associated with selenium deficiency, qPCR was performed on LV tissue to evaluate the expression of cardiac functional markers. When grouped by diet, *Nppb* RNA expression was significantly increased in selenium-deficient mice, whereas *Nppa* and *Gata4* remained unchanged (Figure 2G–I, Supplemental Table 5). When stratified by sex, a marked divergence in molecular responses was found. In male mice, selenium deficiency resulted in a significant upregulation of *Nppb* and *Gata4* expression, whereas *Nppa* showed a nominally significant increase in the unadjusted pairwise comparison that did not remain significant after correction for multiple comparisons (Figure 2J–L, Supplemental Table 5). In contrast, female mice showed no significant increase in these markers. These findings indicate that molecular markers reveal a more pronounced and sex-specific response to selenium deficiency than conventional echocardiographic parameters, highlighting an increased susceptibility of male mice to early cardiac maladaptation.

### 3.3 RNA sequencing reveals expression patterns of cardiac hypertrophy in response to the Selenium-deficient diet in left ventricular tissue

RNA sequencing of isolated LV tissue was performed. For the sequencing effort we used tissues of 10 animals which received control diet (N=5 males and N=5 females) and 10 animals which received the selenium-deficient diet (N=5 males and N=5 females). In summary, we found 2259 genes to be differentially expressed (DEGs; *Padj* <0.05) as the result of selenium deficiency. 1314 (58.2%) of these were found upregulated and 945 (41.8%) were downregulated in mice with selenium deficiency. To reduce potential false positive signals, we applied a cut-off of logFC>|0.5|, reducing the DEGs to a total of 347 (145 upregulated and 202 downregulated) (Table S6).

Within the list of DEGs we found evidence of early cardiac alterations, with the significant increase of Nppb (logFC=0.69; *Padj*=9.11*10-7), Corin (logFC=0.65; *Padj*=4.04*10-5), Tnnt2 (logFC=0.54; *Padj*=0,00057), Ryr2 (logFC=0.53; *Padj*=0,0045) and Myh7 (logFC=0.52; *Padj*=0.012), and a reduction of Tcap (logFC= -0,98; *Padj*=5.3*10-16). This is also reflected by the pathway enrichment analyses that showed significant enrichment for Gene Ontology (GO)-terms related to Cardiac Muscle hypertrophy in response to stress (GO:0014898; *P*=0.02) and Sarcomere organization (GO:0045214; *P*=0.037). As well as significant enrichment for the Monarch (The mammalian phenotype ontology)-term “Cardiac hypertrophy” (MP:0001625; *P*=0.0041). These expression patterns validate the findings observed with echo, pointing towards early systolic dysfunction.

Additionally, among the DEGs we found evidence of selenium deficiency, as reflected by the top significant DEG, Selenow (logFC=-1.20; *Padj*=2.63*10-40), but also the reduction of selenoprotein Selenoh (logFC=-0.74; *Padj*=5.44*10-5). Additionally, the reduction of Sod2 (logFC=-0.51; *Padj*=0.022) exemplifies the reduced antioxidant capacity that is reached as the consequence of reduced selenium availability.

### 3.4 RNA sequencing suggests that the mechanistic relationship between selenium deficiency and HF pathophysiology lies in redox balance

Assessing pathway enrichment analyses in an unbiased manner showed that not only cardiac hypertrophy and sarcomere organization were among the enriched biological processes, but also circadian regulation of gene expression (GO:0032922; *P*=3.43*10-6). Enrichment for circadian rhythm was also observed when assessing KEGG pathway enrichment (mmu04710; *P*= 1.02*10-8). Here 10 members of this pathway were identified as differentially expressed as the consequence of selenium-deficient diet (Clock, Arntl, Npas2, Bhlhe40, Nr1d1, Per3, Cry2, Bhlhe41, Per2 and Skp1a). Of great interest is the apparent overlap in the genes in the pathway “response to redox state” (GO:0051775; *P*=0.0056; Clock, Arntl, Npas2 and Ryr2), which are also found in “circadian regulation of gene expression” and “cardiac muscle hypertrophy”.

Looking at enrichment for cellular component Go-terms we again found enrichment for circadian alterations (GO:1990513; *P*=0.0069) and cellular components specific for cardiac tissue (Z-disc, GO:0030018; *P*=0.012 and Sarcomere, GO:0030017; *P*=0.0018). Additionally, mitochondrial membrane related genes were found enriched, including mitochondrial Sod (Sod2; logFC=-0.52; *Padj*=0.022), which is a major component of the metabolic machinery that handles reactive oxygen species (ROS) in the mitochondria.

### 3.5 Sex specificity of differential transcriptomics suggests males to be more sensitive to selenium deficiency induced detrimental changes

The PCA plot obtained based on the variance-stabilized RNA sequencing data showed a distinct separation, not only based on diet, but also based on sex (Figure 4A). Stratifying the transcriptomics data by sex showed some marked differences. Nonetheless, both sexes showed a clear sign of selenium deprivation with Selenow being the most significantly downregulated transcript in both male and female mice (logFC=-1.61; *Padj*= 6.77*10-72 and logFC=-1.04; *Padj*= 1.86*10-35, respectively). Assessing the transcriptional differences between sex in the deficiency group shows a marked enrichment for GO-term ‘biological processes’ involved in muscle contraction (GO:0006936; *P*=3.36*10-8; eg. ATP2a2, Myh6, Ryr2, TnnT2 and Myh7), with Nppb significantly upregulated in male LV tissue in the Se-deficient group (logFC=1.03; *Padj*= 0,00016), but unchanged in females (Figure 4B). Stratifying GO-term processes by sex shows that males primarily show an enrichment of cytoplasmic translation (GO:0002181; *P*=7.35*10-46) (Figure 4C). In females, we find mostly enrichment for oxidative phosphorylation (GO:0006119; *P*=2.18*10-29) and other mitochondrial related processes (Figure 4D). Notably, for both sexes, an enrichment for circadian rhythm was observed among GO and KEGG terms with distinct differences in transcriptional regulation (Figure 4E-F).

**Figure 4:**
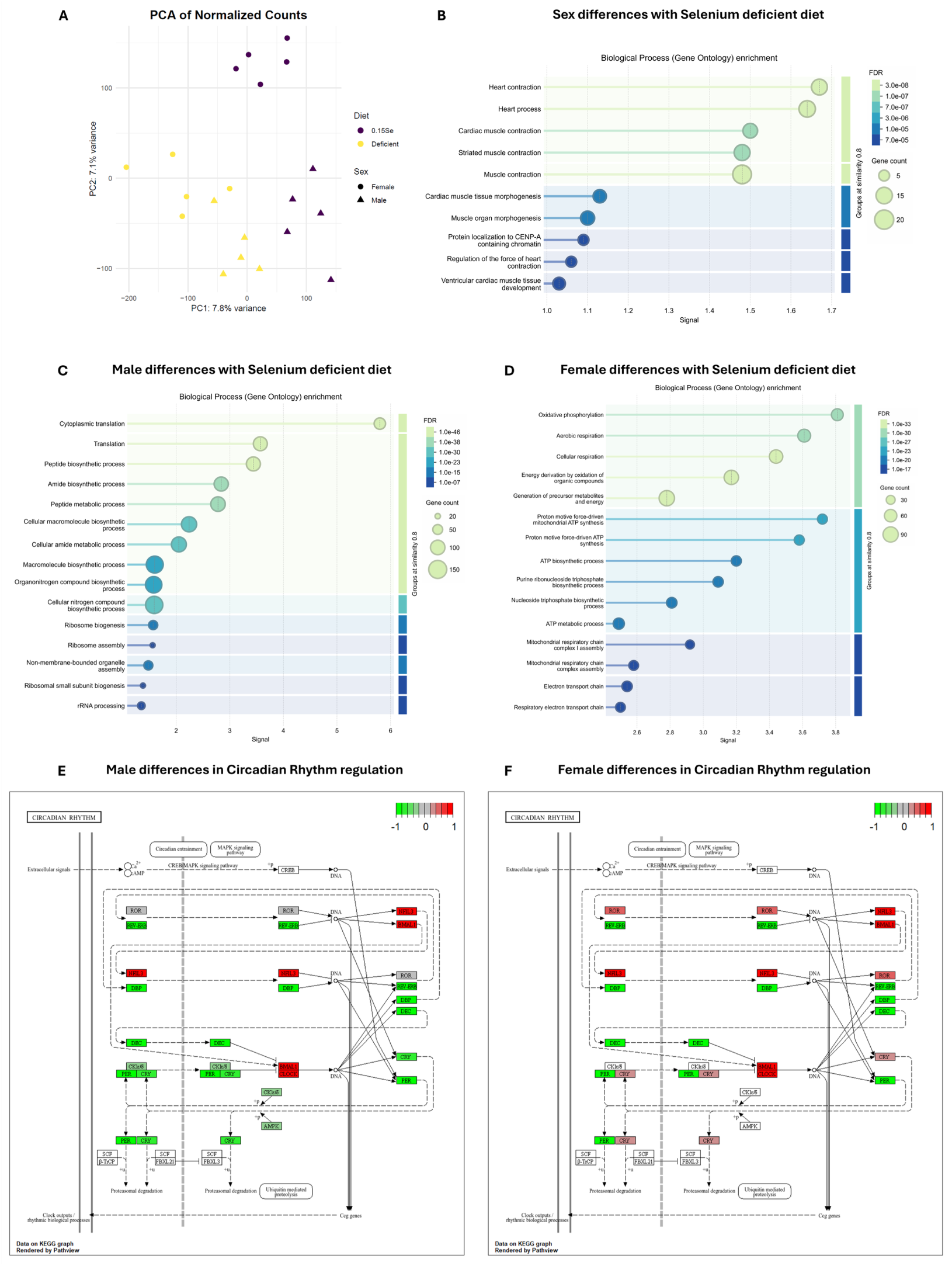
RNA sequencing analyses of left ventricle tissue of selenium-sufficient and selenium-deficient mice. **A)** PCA plot. **B)** GO-term enrichment for Biological Processes involved in sex-based differences with selenium-deficient diet. **C)** GO-term enrichment for Biological Processes based on male differences with selenium-deficient diet. **D)** GO-term enrichment for Biological Processes based on female differences with selenium-deficient diet. **E)** KEGG pathway visualization with a colour map indicating the direction of expression changes in males for the circadian rhythm pathway. F) KEGG pathway visualization with a colour map indicating the direction of expression changes in females for the circadian rhythm pathway. For sex-stratified RNA-sequencing analyses, n = 5 mice per sex per dietary group (independent biological replicates). Differential expression was analysed using DESeq2 with Wald tests and Benjamini–Hochberg correction for multiple testing. GO-term enrichment was performed using clusterProfiler::enrichGO() with hypergeometric over-representation analysis and Benjamini–Hochberg correction; q values < 0.05 were considered significant. KEGG pathway enrichment was performed using clusterProfiler::enrichKEGG() with Benjamini–Hochberg correction and a q-value cutoff of 0.05. Enriched KEGG pathways were visualized using Pathview by mapping gene-level log2 fold-change values onto the KEGG pathway diagrams.

### 3.6 Sex specificity underlying the mechanistic relationship between selenium deficiency and HF pathophysiology points to differential redox balance

In order to explain the predominant development of early systolic dysfunction in males, we assessed molecular pathways directly involved by selenium deficiency. Selenium availability is directly involved in cellular ROS response and redox balance through selenoproteins GPX1 and GPX4, which reduce hydrogen peroxide to water by using GSH. Both proteins are upregulated most prominently in deficient males compared to deficient female mice (Figure 5B-C), indicating an increase in ROS. This was accompanied by a lower GSH/GSSG ratio in the LV tissues of males in the selenium deficient group (Figure 5D-E), reflecting higher hydrogen peroxide reduction and overall poor redox balance. As the increased GPX protein levels indicate a higher presence of oxidative stress, we assessed SOD2 and Catalase protein levels (Figure 5A). Catalase protein expression showed a significant effect of sex (*P*=0.024), but not of diet (*P=*0.059) (Figure 5F). Expression of SOD2, showed a trend of upregulation in selenium deficiency (*P*=0.062). The upregulated oxidative stress response proteins and affected GSH/GSSG ratio indicate a poor redox state specific for selenium deficient males. As such, the involved NAD+/NADH showed a sex-specific larger ratio value in female mice LV compared to male mice LV (*P*=0.019), but no effect of diet (Figure 5H-I).

**Figure 5:**
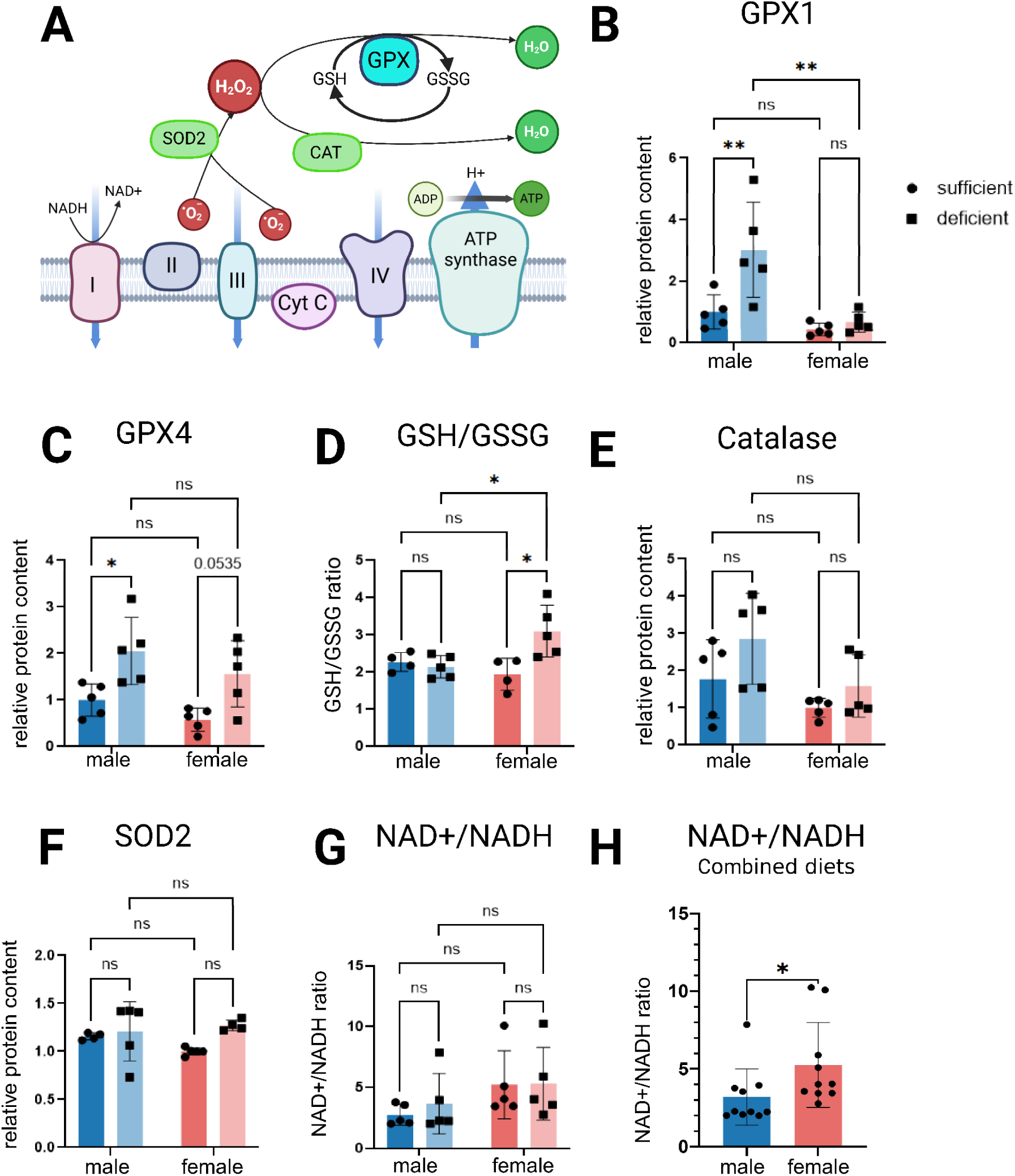
Reactive oxygen species pathway and redox balance in selenium-deficient mouse left ventricular tissue. **A)** Schematic overview of the reactive oxygen species response pathway in the mitochondrial electron transport chain. **B–G)** GPX1 protein levels (B), GPX4 protein levels (C), GSH/GSSG ratio (D), catalase protein levels (E), SOD2 protein levels (F), and NAD+/NADH ratio (G), stratified by sex and selenium-sufficient or selenium-deficient diet. **H)** NAD+/NADH ratio in male versus female left ventricular tissue. For panels B–G, n = 5 independent biological replicates per group. Outliers were identified using the ROUT method (Q = 1%). Data were analysed using two-way ANOVA followed by Tukey’s multiple-comparisons test. For panel H, the two male groups were combined (n = 10 independent biological replicates) and the two female groups were combined (n = 10 independent biological replicates), and the groups were compared using the Mann–Whitney U test. *P < 0.05, **P < 0.01.

### 3.7 Redox imbalance affects sex-dependent dysregulation of Circadian Rhythm

The sex dependent NAD+/NADH balance in combination with the pathway enrichment for circadian rhythm factors rationalized further investigation of the effects of selenium deficiency on the circadian rhythm pathway in mice LV due to the regulatory role of NAD+ on SIRT1 in the circadian rhythm (Figure 6A). The deacetylase SIRT1 protein is specifically upregulated in male selenium deficient mice LV, with no changes in female mice (Figure 6B). SIRT1 activity, as demonstrated by deacetylation of FOXO1 (Figure 6C&E), is higher in selenium deficient males compared to selenium deficient females. Sex specific circadian rhythm deregulation is also confirmed by PER2 protein levels, as female mice LV tissue shows a significantly lower amount of PER2 (Figure 6D&F). This sex-specific regulatory difference downstream of selenium deficiency is in line with the male-prominent heart failure progression.

**Figure 6:**
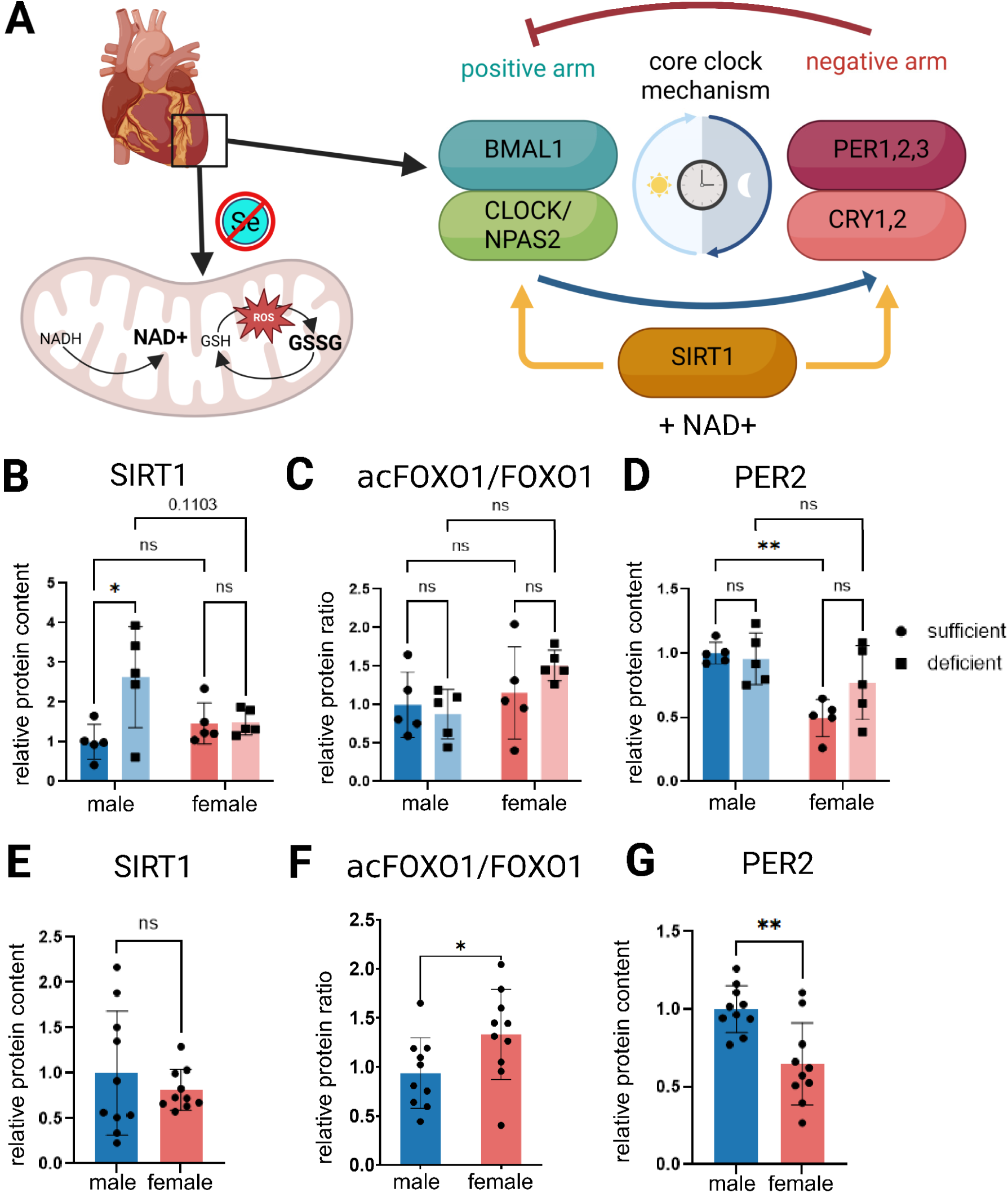
Circadian rhythm pathway regulation assessment. **A)** Schematic of the hypothesis that selenium deficiency in the left ventricle alters redox state and circadian rhythm. **B)** SIRT1 protein levels stratified by sex and selenium-sufficient or selenium-deficient diet. **C)** Acetyl-FOXO1/FOXO1 protein level ratio stratified by sex and selenium-sufficient or selenium-deficient diet. **D)** PER2 protein levels stratified by sex and selenium-sufficient or selenium-deficient diet. **E)** Relative SIRT1 protein level in male versus female left ventricular tissue, normalized to the male group. **F)** Acetyl-FOXO1/FOXO1 protein level ratio in male versus female left ventricular tissue. **G)** Relative PER2 protein level in male versus female left ventricular tissue, normalized to the male group. For panels B– D, n = 5 independent biological replicates per group. Data were analysed using two-way ANOVA followed by Tukey’s multiple-comparisons test. For panels E–G, the two male dietary groups were combined (n = 10 independent biological replicates) and the two female dietary groups were combined (n = 10 independent biological replicates), and males and females were compared using an unpaired two-tailed Student’s t-test. *P < 0.05, **P < 0.01.

## 4. DISCUSSION

In this study, we show that dietary selenium deficiency induces early, sexual dimorphic subclinical cardiac dysfunction characterized by impaired myocardial deformation, activation of hypertrophic gene programmes, and disruption of redox and circadian homeostasis. Building upon our previous findings that selenium depletion exacerbates heart failure progression [16], induces mitochondrial oxidative stress in cardiomyocytes[15] and impairs immunoregulatory processes [24], we now demonstrate that even in the absence of overt disease, dietary selenium deficiency initiates early, subclinical cardiac maladaptation.

Although ejection fraction and fractional shortening were preserved after 12 weeks, GLS was significantly reduced, indicating early systolic dysfunction not detected by conventional echocardiography. This was accompanied by increased LVPW, consistent with early structural remodelling. These findings show GLS as a sensitive marker of micronutrient-induced cardiac dysfunction preceding overt functional decline. This was reflected by molecular evidence of early cardiac stress. Selenium-deficient mice showed increased Nppb expression, especially in males. This sex-specific response was further supported by male-specific upregulation of Nppa and Gata4, indicating activation of hypertrophic transcriptional programmes, whereas females showed minimal induction despite similar systemic selenium reduction. Sexual dimorphism in selenium deficiency has been observed before regarding metabolic syndrome, type 2 diabetes, hypertension and prevalence of heart disease[25, 26].

Consistent with the established role of selenium in antioxidant defence and redox balance [14, 27, 28], we observed marked reductions in plasma selenium levels and molecular signatures of selenium depletion in LV tissues. Selenoproteins (e.g. SelenoW) were downregulated, confirming systemic deficiency. Importantly, these effects occurred independently of other trace element alterations, apart from an increase in plasma copper, which may further exacerbate redox imbalance[29]. RNA sequencing of LV tissue revealed a distinct pattern of DEGs consistent with early maladaptive cardiac remodelling. Upregulation of Nppb, Corin, Tnnt2, Ryr2, and Myh7, together with the downregulation of Tcap, suggest that selenium deficiency disrupts sarcomeric organization and contractile function, corroborated by pathway enrichment for cardiac muscle hypertrophy and sarcomere organization.

A central finding of this study is the identification of redox imbalance as a key mediator of these effects. Selenium deficiency reduced antioxidant capacity and increased oxidative stress, consistent with prior work linking it to ROS accumulation and mitochondrial dysfunction [15]. This was more pronounced in males, which showed increased GPX1/GPX4 protein levels, reduced GSH/GSSG ratios, and an unfavourable NAD⁺/NADH balance, indicating insufficient compensatory antioxidant responses. Enhanced FOXO1 deacetylation further supports activation of antioxidant signalling via SIRT1-mediated pathways[30]. In contrast, females demonstrated a more favourable NAD⁺/NADH ratio and enrichment of mitochondrial and oxidative phosphorylation pathways, suggesting a more adaptive metabolic response.

Notably, our data identified redox imbalance and circadian rhythm dysregulation as a downstream consequence of selenium deficiency. Both transcriptomic and pathway analyses revealed robust enrichment of circadian gene networks, including altered expression of core clock components such as Clock, Arntl, Npas2, Nr1d1, Cry2, and Per2. The overlap between circadian and redox-responsive pathways underscores the tight coupling between metabolic state and circadian regulation in the heart, consistent with previous work demonstrating that disruption of circadian control increases cardiac vulnerability through maladaptive redox signalling[31]. Moreover, circadian misalignment has been implicated in cardiomyopathy, metabolic dysfunction, and impaired immune surveillance [32], supporting the notion that selenium deficiency may act as an upstream modifier of circadian regulation.

Mechanistically, this likely involves NAD⁺-dependent SIRT1, linking redox status to circadian control[33]. Increased SIRT1 activity in selenium-deficient males, reflected by FOXO1 deacetylation, likely reflects altered NAD⁺/NADH balance. SIRT1 modulates core clock machinery and thereby alters the temporal regulation of cardiac gene expression[34, 35], providing a mechanistic bridge between redox imbalance and circadian reprogramming.

Consistent with this, we observed sex-specific alterations in PER2 protein levels, a key negative regulator of the circadian clock. Female mice exhibited reduced PER2 protein levels, whereas males did not, despite increased SIRT1 activity. This suggests that selenium deficiency induces distinct circadian adaptations in males and females. In males, enhanced SIRT1–FOXO1 signalling may drive maladaptive circadian reprogramming without reducing PER2 abundance, whereas in females, reduced PER2 levels may reflect a dampened circadian amplitude that could be protective under conditions of metabolic stress. These findings align with previous reports demonstrating sex-dependent differences in circadian regulation and cardiac outcomes, including increased susceptibility to circadian disruption–induced cardiomyopathy in males[36] and differential circadian responses to physiological stress[37].

Overall, selenium deficiency disrupts cardiac redox balance, activating stress-responsive and hypertrophic pathways, and reprogramming of circadian gene expression [38]. A reference dataset of rhythmic genes[39], reflects the strong enrichment among DEGs, indicating broad circadian output remodelling. Females showed greater involvement of circadian rhythm genes, which may reflect adaptive circadian engagement consistent with preserved function and redox balance. This supports a redox–circadian axis underlying sex-dependent susceptibility.

From a translational perspective, these findings suggest that subclinical selenium deficiency may contribute to early myocardial dysfunction not detectable by conventional systolic measures. They also emphasize sex-specific vulnerability and identify the redox–circadian interface, including SIRT1 signalling, as a potential therapeutic target in selenium-associated cardiac dysfunction.

## CONCLUSION

This study demonstrates that selenium deficiency induces molecular and functional cardiac alterations, characterized by early remodelling, impaired myocardial deformation, mitochondrial stress, and circadian gene reprogramming. These coordinated changes highlight selenium’s essential role in maintaining cardiac redox and circadian homeostasis. These findings expand our understanding of selenium’s systemic protective functions and emphasize the complex interplay between micronutrient status, redox-circadian biology, and cardiac health. Given the increasing recognition of subclinical selenium deficiency in human populations, particularly in regions with low dietary selenium availability, these findings have important translational relevance for cardiovascular risk assessment. Our data provide a foundation for nutritional interventions targeting selenium deficiency that may restore cardiac function through both direct and indirect redox-sensitive pathways.

## SUPPLEMENTARY MATERIAL

## Acknowledgements

We gratefully acknowledge Renate Jagersma, Andriana Papadaki and Martin M. Dokter for their valuable assistance with experimental procedures, sample processing, and data acquisition. We also thank the staff of the UMCG Central Animal Facility (CDP) for their support, in particular Daryll S. Eichhorn, Michel Weij, and Miriam van der Meulen.

## Author contributions

N.B. and P.v.d.M. supervised the study, contributed to study design and data interpretation, and critically revised the manuscript. X.G. designed the study, led the experimental work, performed most experiments, analysed and interpreted the data, prepared the figures, and drafted the manuscript. F.L. contributed to additional experiments, data interpretation, and manuscript drafting. L.G. contributed to data analysis, downstream visualization and interpretation. J.Z. S.Z., A.M.F, and E.M.S. contributed to animal experimental procedures. H.H.W.S. contributed to laboratory coordination, project administration, and regulatory oversight. L.S. provided expertise, resources, and supervision related to selenium biology and trace element analysis. All authors reviewed and approved the final manuscript. *F.L. and L.G. contributed equally to this work*.

## Data availability

All data that support the findings of this study are available from the corresponding author upon reasonable request.

## References

1. Heidenreich PA, Bozkurt B, Aguilar D, Allen LA, Byun JJ, Colvin MM, Deswal A, Drazner MH, Dunlay SM, Evers LR et al: 2022 AHA/ACC/HFSA Guideline for the Management of Heart Failure: A Report of the American College of Cardiology/American Heart Association Joint Committee on Clinical Practice Guidelines. Circulation 2022, 145(18):e895–e1032.

2. Ziaeian B, Fonarow GC: Epidemiology and aetiology of heart failure. Nat Rev Cardiol 2016, 13(6):368–378.

3. Al-Mubarak AA, van der Meer P, Bomer N: Selenium, Selenoproteins, and Heart Failure: Current Knowledge and Future Perspective. Curr Heart Fail Rep 2021, 18(3):122–131.

4. Loscalzo J: Keshan disease, selenium deficiency, and the selenoproteome. N Engl J Med 2014, 370(18):1756–1760.

5. Klip IT, Comin-Colet J, Voors AA, Ponikowski P, Enjuanes C, Banasiak W, Lok DJ, Rosentryt P, Torrens A, Polonski L et al: Iron deficiency in chronic heart failure: an international pooled analysis. Am Heart J 2013, 165(4):575–582.e573.

6. Yoshihisa A, Abe S, Kiko T, Kimishima Y, Sato Y, Watanabe S, Kanno Y, Miyata-Tatsumi M, Misaka T, Sato T et al: Association of Serum Zinc Level With Prognosis in Patients With Heart Failure. J Card Fail 2018, 24(6):375–383.

7. Roberts CG, Ladenson PW: Hypothyroidism. Lancet 2004, 363(9411):793–803.

8. Hughes CM, Woodside JV, McGartland C, Roberts MJ, Nicholls DP, McKeown PP: Nutritional intake and oxidative stress in chronic heart failure. Nutr Metab Cardiovasc Dis 2012, 22(4):376–382.

9. McKeag NA, McKinley MC, Harbinson MT, McGinty A, Neville CE, Woodside JV, McKeown PP: Dietary Micronutrient Intake and Micronutrient Status in Patients With Chronic Stable Heart Failure: An Observational Study. J Cardiovasc Nurs 2017, 32(2):148–155.

10. Fairweather-Tait SJ, Bao Y, Broadley MR, Collings R, Ford D, Hesketh JE, Hurst R: Selenium in human health and disease. Antioxid Redox Signal 2011, 14(7):1337–1383.

11. Cheng YY, Qian PC: The effect of selenium-fortified table salt in the prevention of Keshan disease on a population of 1.05 million. Biomed Environ Sci 1990, 3(4):422–428.

12. Li Y, Peng T, Yang Y, Niu C, Archard LC, Zhang H: High prevalence of enteroviral genomic sequences in myocardium from cases of endemic cardiomyopathy (Keshan disease) in China. Heart 2000, 83(6):696–701.

13. Guillin OM, Vindry C, Ohlmann T, Chavatte L: Selenium, Selenoproteins and Viral Infection. Nutrients 2019, 11(9).

14. Rayman MP: Selenium and human health. Lancet 2012, 379(9822):1256–1268.

15. Bomer N, Grote Beverborg N, Hoes MF, Streng KW, Vermeer M, Dokter MM, J IJ, Anker SD, Cleland JGF, Hillege HL et al: Selenium and outcome in heart failure. Eur J Heart Fail 2020, 22(8):1415–1423.

16. Al-Mubarak AA, Grote Beverborg N, Suthahar N, Gansevoort RT, Bakker SJL, Touw DJ, de Boer RA, van der Meer P, Bomer N: High selenium levels associate with reduced risk of mortality and new-onset heart failure: data from PREVEND. Eur J Heart Fail 2022, 24(2):299–307.

17. Percie du Sert N, Hurst V, Ahluwalia A, Alam S, Avey MT, Baker M, Browne WJ, Clark A, Cuthill IC, Dirnagl U et al: The ARRIVE guidelines 2.0: updated guidelines for reporting animal research. J Physiol 2020, 598(18):3793–3801.

18. Stege NM, Oliveira Nunes Teixeira V, Zijlstra SN, Feringa AM, de Boer RA, Silljé HHW: Deletion of DWORF does not affect cardiac function in aging and in PLN-R14del cardiomyopathy. Am J Physiol Heart Circ Physiol 2024, 326(3):H870–h876.

19. Eijgenraam TR, Stege NM, Oliveira Nunes Teixeira V, de Brouwer R, Schouten EM, Grote Beverborg N, Sun L, Später D, Knöll R, Hansson KM et al: Antisense Therapy Attenuates Phospholamban p.(Arg14del) Cardiomyopathy in Mice and Reverses Protein Aggregation. Int J Mol Sci 2022, 23(5).

20. Moghaddam A, Heller RA, Sun Q, Seelig J, Cherkezov A, Seibert L, Hackler J, Seemann P, Diegmann J, Pilz M et al: Selenium Deficiency Is Associated with Mortality Risk from COVID-19. Nutrients 2020, 12(7).

21. Stosnach H: Environmental trace-element analysis using a benchtop total reflection X-ray fluorescence spectrometer. Anal Sci 2005, 21(7):873–876.

22. Eijgenraam TR, Boogerd CJ, Stege NM, Oliveira Nunes Teixeira V, Dokter MM, Schmidt LE, Yin X, Theofilatos K, Mayr M, van der Meer P et al: Protein Aggregation Is an Early Manifestation of Phospholamban p.(Arg14del)-Related Cardiomyopathy: Development of PLN-R14del-Related Cardiomyopathy. Circ Heart Fail 2021, 14(11):e008532.

23. Stege NM, Eijgenraam TR, Oliveira Nunes Teixeira V, Feringa AM, Schouten EM, Kuster DWD, van der Velden J, Wolters AHG, Giepmans BNG, Makarewich CA et al: DWORF Extends Life Span in a PLN-R14del Cardiomyopathy Mouse Model by Reducing Abnormal Sarcoplasmic Reticulum Clusters. Circ Res 2023, 133(12):1006–1021.

24. Al-Mubarak AA, Markousis Mavrogenis G, Guo X, De Bruyn M, Nath M, Romaine SPR, Grote Beverborg N, Arevalo Gomez K, Zijlstra SN, van Veldhuisen DJ et al: Biomarker and transcriptomics profiles of serum selenium concentrations in patients with heart failure are associated with immunoregulatory processes. Redox Biol 2024, 70:103046.

25. Weening EH, Al-Mubarak AA, Dokter MM, Dickstein K, Lang CC, Ng LL, Metra M, van Veldhuisen DJ, Touw DJ, de Boer RA et al: Sexual dimorphism in selenium deficiency is associated with metabolic syndrome and prevalence of heart disease. Cardiovascular Diabetology 2023, 22(1):8.

26. Demircan K, Hybsier S, Chillon TS, Vetter VM, Rijntjes E, Demuth I, Schomburg L: Sex-specific associations of serum selenium and selenoprotein P with type 2 diabetes mellitus and hypertension in the Berlin Aging Study II. Redox Biol 2023, 65:102823.

27. Alehagen U, Johansson P, Björnstedt M, Rosén A, Dahlström U: Cardiovascular mortality and N-terminal-proBNP reduced after combined selenium and coenzyme Q10 supplementation: a 5-year prospective randomized double-blind placebo-controlled trial among elderly Swedish citizens. Int J Cardiol 2013, 167(5):1860–1866.

28. Johansson P, Dahlström Ö, Dahlström U, Alehagen U: Effect of selenium and Q10 on the cardiac biomarker NT-proBNP. Scand Cardiovasc J 2013, 47(5):281–288.

29. Linder MC: Ceruloplasmin and other copper binding components of blood plasma and their functions: an update. Metallomics 2016, 8(9):887–905.

30. Akasaki Y, Alvarez-Garcia O, Saito M, Caramés B, Iwamoto Y, Lotz MK: FoxO transcription factors support oxidative stress resistance in human chondrocytes. Arthritis Rheumatol 2014, 66(12):3349–3358.

31. Lecacheur M, Ammerlaan DJM, Dierickx P: Circadian rhythms in cardiovascular (dys)function: approaches for future therapeutics. npj Cardiovascular Health 2024, 1(1):21.

32. Durgan DJ, Young ME: The cardiomyocyte circadian clock: emerging roles in health and disease. Circ Res 2010, 106(4):647–658.

33. Putker M, O’Neill JS: Reciprocal Control of the Circadian Clock and Cellular Redox State - a Critical Appraisal. Mol Cells 2016, 39(1):6–19.

34. Kawashima T, Inuzuka Y, Okuda J, Kato T, Niizuma S, Tamaki Y, Iwanaga Y, Kawamoto A, Narazaki M, Matsuda T et al: Constitutive SIRT1 overexpression impairs mitochondria and reduces cardiac function in mice. J Mol Cell Cardiol 2011, 51(6):1026–1036.

35. Alcendor RR, Gao S, Zhai P, Zablocki D, Holle E, Yu X, Tian B, Wagner T, Vatner SF, Sadoshima J: Sirt1 regulates aging and resistance to oxidative stress in the heart. Circ Res 2007, 100(10):1512–1521.

36. Alibhai FJ, Reitz CJ, Peppler WT, Basu P, Sheppard P, Choleris E, Bakovic M, Martino TA: Female ClockΔ19/Δ19 mice are protected from the development of age-dependent cardiomyopathy. Cardiovasc Res 2018, 114(2):259–271.

37. Anderson ST, Meng H, Brooks TG, Tang SY, Lordan R, Sengupta A, Nayak S, Mřela A, Sarantopoulou D, Lahens NF et al: Sexual dimorphism in the response to chronic circadian misalignment on a high-fat diet. Sci Transl Med 2023, 15(696):eabo2022.

38. Das MK, De Ryck E, Jorgensen IL, Zienolddiny-Narui S, Erdem JS: Circadian rhythm disruption in cardiovascular disease: a systematic review and meta-analysis of mechanistic evidence from animal models. BMC Med 2026, 24(1):73.

39. Mure LS, Le HD, Benegiamo G, Chang MW, Rios L, Jillani N, Ngotho M, Kariuki T, Dkhissi-Benyahya O, Cooper HM et al: Diurnal transcriptome atlas of a primate across major neural and peripheral tissues. Science 2018, 359(6381).

