## Supplemental Figures for "Dietary selenium deficiency drives sex-specific circadian disturbance through redox imbalance and causes early systolic dysfunction in mice"

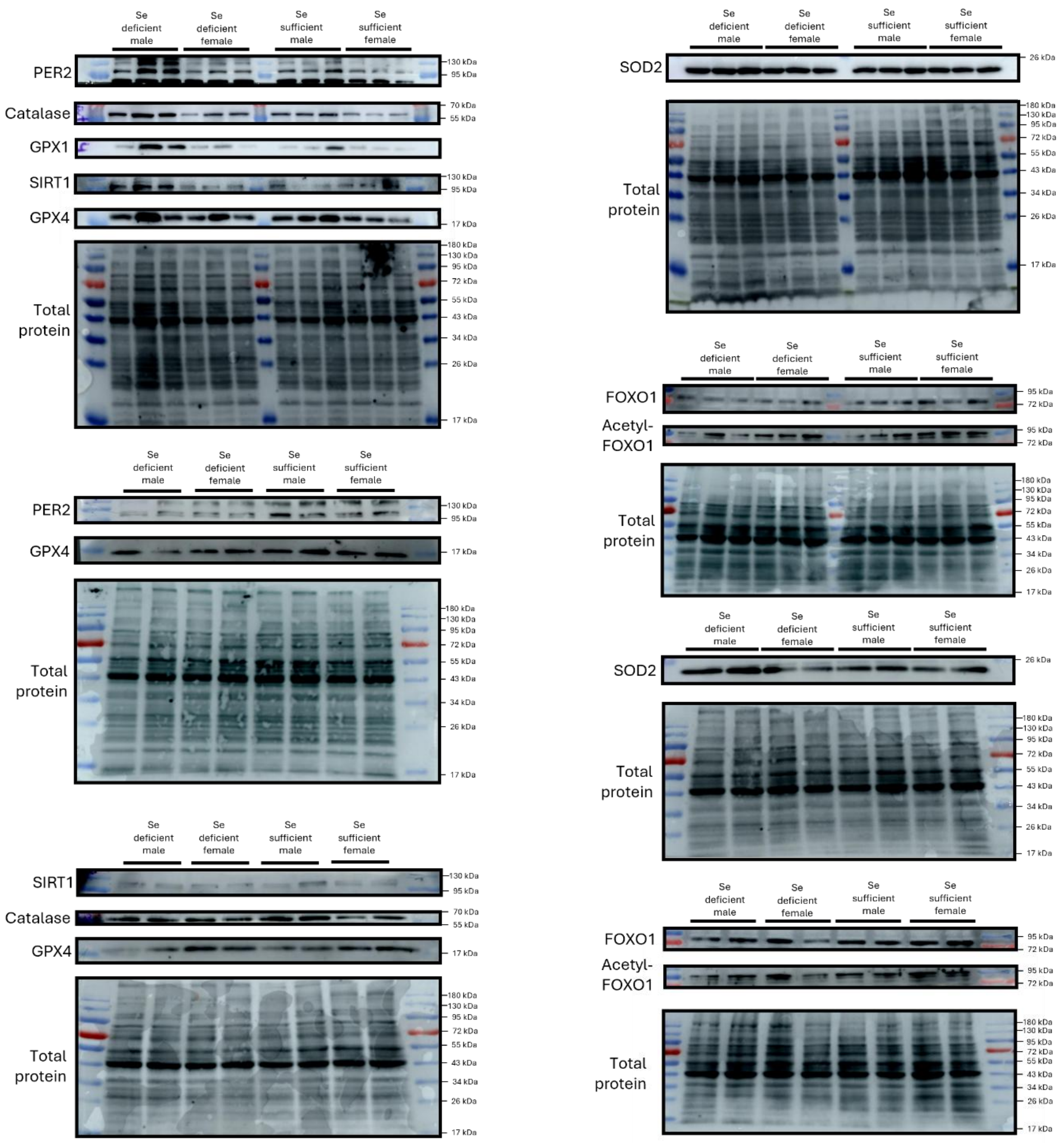


**Supplementary Figure 1. Representative Western blot images of redox- and circadian-related proteins in left ventricular tissue.** Representative Western blot images are shown for proteins involved in circadian regulation, antioxidant defence, and redox-sensitive signalling, including PER2, catalase, GPX1, SIRT1, GPX4, SOD2, total FOXO1, and acetylated FOXO1. PER2 was assessed as a negative-arm factor of the circadian rhythm pathway, while SIRT1 was assessed as an NAD+-dependent deacetylase and circadian rhythm regulator. Catalase, GPX1 (glutathione peroxidase 1), GPX4 (glutathione peroxidase 4), and SOD2 (superoxide dismutase 2) were assessed as antioxidant defence proteins, and FOXO1 acetylation was assessed as a marker of acetylation-regulated oxidative stress-responsive transcriptional regulation. n = 5 independent biological replicates per group.

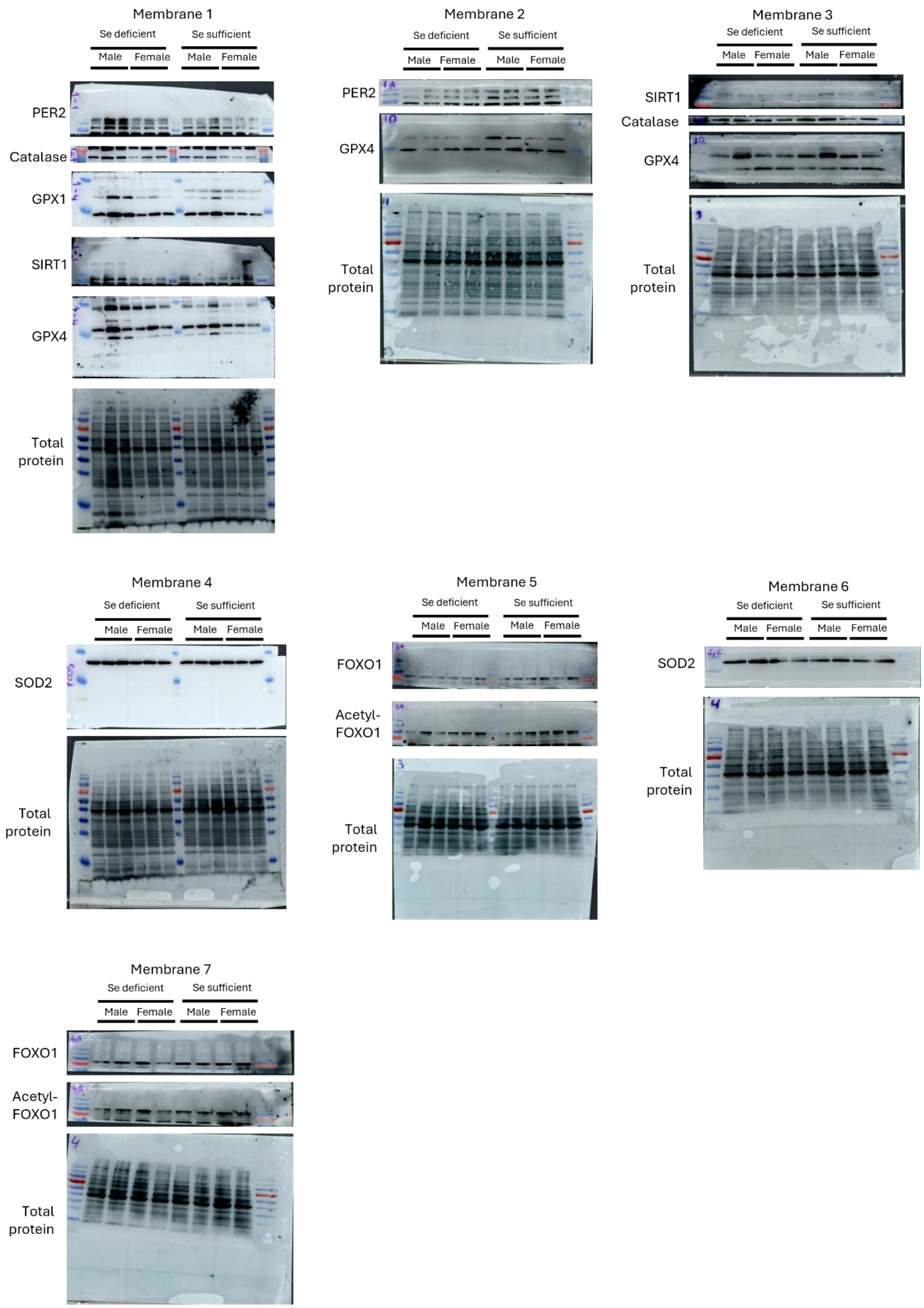

Membrane 1
Se deficient
Se sufficient
Male
Female
PER2
Catalase
GPX1
SIRT1
GPX4
Total protein
Membrane 2
Se deficient
Se sufficient
Male
Female
PER2
GPX4
Total protein
Membrane 3
Se deficient
Se sufficient
Male
Female
SIRT1
Catalase
GPX4
Total protein
Membrane 4
Se deficient
Se sufficient
Male
Female
SOD2
Total protein
Membrane 5
Se deficient
Se sufficient
Male
Female
FOXO1
Acetyl-FOXO1
Total protein
Membrane 6
Se deficient
Se sufficient
Male
Female
SOD2
Total protein
Membrane 7
Se deficient
Se sufficient
Male
Female
FOXO1
Acetyl-FOXO1
Total protein

**Supplementary figures 2: Uncropped Western blot images of redox- and circadian-related proteins in left ventricular tissue.**

Uncropped Western blot images corresponding to the representative blots shown in Supplementary Figure 1 are provided for PER2, catalase, GPX1, SIRT1, GPX4, SOD2, total FOXO1, and acetylated FOXO1. These proteins were analyzed to assess circadian regulation, antioxidant defence, and redox-sensitive signalling in left ventricular tissue. n = 5 independent biological replicates per group. Each blot contained either two biological replicates (membranes 2, 3, 6, and 7) or three biological replicates (membranes 1, 4, and 5)
