## Supplemental Tables for "Dietary selenium deficiency drives sex-specific circadian disturbance through redox imbalance and causes early systolic dysfunction in mice"

**Index supplementary tables.**

**Supplemental table 1.** Plasma elements concentration.

**Supplemental table 2.** Body weight increase and organ weights normalized to tibia length

**Supplemental table 3.** Fibrosis area in left ventricle and major organs(%) .

**Supplemental table 4.** List of primers.

**Supplemental table 5**. Left ventricular tissue RNA expression(fold change, corrected by 36B4)

**Supplemental table 6.** RNA sequencing reveals expression patterns of cardiac hypertrophy in response to the Selenium-deficient diet in left ventricular tissue (**separate .xls file**).

**Supplemental table 7.** List Antibodies.

**Supplemental table 1. Plasma elements concentrations**

|  | **Se-0.15 ppm (control)** | | **Se-deficient** | |
| --- | --- | --- | --- | --- |
| **Elements** | **Males, n=10** | **Females, n=9** | **Males, n=9** | **Females, n=9** |
| Selenium (µg/l) | 352.1±14.84 | 457±15.98 | 51.47±2.26^****^ | 68.31±3.40^####^ |
|  | **Males, n=10** | **Females, n=9** | **Males, n=9** | **Females, n=10** |
| Calcium (mg/l) | 101.3±8.55 | 101.8±2.84 | 99.92±3.76 | 98.43±3.29 |
| Iron (µg/l) | 3982±269.30 | 3563±244.40 | 5089±658.20 | 3712±395.60 |
| Copper (µg/l) | 564.3±27.26 | 696.9±17.13 | 560.7±19.26 | 561.6±16.56^###^ |
| Zinc (µg/l) | 1061±27.21 | 1119±39.04 | 1098±31.52 | 1144±33.59 |

**Supplemental table 1. Plasma elements concentrations.** Data are presented as mean ± SEM. For each element, data were analysed using two-way ANOVA followed by Šídák’s multiple-comparisons test. Asterisks indicate comparisons between selenium-deficient and selenium-sufficient males, and hash symbols indicate comparisons between selenium-deficient and selenium-sufficient females. ###P < 0.001; ****P < 0.0001; ####P < 0.0001.

**Supplemental table 2. Body weight increase and organ weights normalized to tibia length**

|  | **Se-0.15 ppm (control)** | | **Se-deficient** | |
| --- | --- | --- | --- | --- |
|  | **Males, n=10** | **Females, n=10** | **Males, n=9** | **Females, n=10** |
| Body weight increase(g) | 6.35±0.88 | 3.45±0.25 | 9.26±0.80^**^ | 3.10±0.31 |
|  | **n=4** | **n=5** | **n=5** | **n=4** |
| Heart weight/tibia length (mg/mm) | 8.64±0.77 | 6.79±0.25 | 7.80±0.34 | 6.35±0.25 |
| Liver/tibia length (mg/mm) | 73.36±5.17 | 62.28±2.90 | 80.66±4.34 | 59.26±1.75 |
| Kidney/tibia length (mg/mm) | 10.25±0.43 | 7.95±0.30 | 10.76±0.53 | 7.22±0.49 |
| Spleen/tibia length (mg/mm) | 4.90±0.13 | 6.34±0.29 | 4.48±0.13 | 4.59±0.15^###^ |

**Supplemental Table 2.** **Body weight increase and organ weights normalized to tibia length.** Data are presented as mean ± SEM. For each parameter, data were analysed using two-way ANOVA followed by Šídák’s multiple-comparisons test. Asterisks indicate comparisons between selenium-deficient and selenium-sufficient males, and hash symbols indicate comparisons between selenium-deficient and selenium-sufficient females. ***P < 0.001; ####P < 0.0001.

**Supplemental Table 3. Fibrosis area in left ventricle and major organs（%）**

|  | **Se-0.15 ppm (control)** | | **Se-deficient** | |
| --- | --- | --- | --- | --- |
| **Organs** | **Males, n=5** | **Females, n=5** | **Males, n=5** | **Females, n=5** |
| Left ventricle | 0.4540 ± 0.06 | 0.3596 ± 0.03 | 0.3440 ± 0.03 | 0.3925 ± 0.05 |
| Kidney | 1.874 ± 0.25 | 4.765 ± 0.38 | 1.102 ±0.31 | 1.890 ± 0.44^###^ |
| Spleen | 0.7080 ± 0.19 | 1.462 ± 0.34 | 1.424 ± 0.25 | 0.9687 ± 0.15 |
| Liver | 0.4950 ± 0.03 | 1.001 ± 0.10 | 0.5688 ± 0.05 | 0.6292 ± 0.09^#^ |
| Muscle | 4.360 ± 0.37 | 4.624 ± 1.28 | 2.596 ± 0.53 | 4.341 ± 0.76 |

**Supplemental Table 3. Fibrosis area in left ventricle and major organs.** Data are presented as mean ± SEM. Data were analysed using two-way ANOVA followed by Šídák’s multiple-comparisons test. Asterisks indicate comparisons between selenium-deficient and selenium-sufficient males, and hash symbols indicate comparisons between selenium-deficient and selenium-sufficient females. #P < 0.05; ###P < 0.001.

**Supplemental table 4. List of primers**

| **Gene** | **Sequence** |
| --- | --- |
| 36B4 | Fwd: AAGCGCGTCCTGGCATTGTC Rev: GCAGCCGCAAATGCAGATGG |
| Nppa | Fwd: GCTTCCAGGCCATATTGGAG Rev: GGTGGTCTAGCAGGTTCTTG |
| Nppb | Fwd: CTCCTATCCTCTGGGAAGTC Rev: CCGATCCGGTCTATCTTGTG |
| GATA4 | Fwd: AACGGAAGCCCAAGAACCTG Rev: GTGGCATTGCTGGAGTTACC |
| GPX1 | Fwd: CTCTTCATTCTTGCCATTCTCCTG Rev: ACAGTCCACCGTGTATGCCTTC |
| GPX2 | Fwd: GTTCTCGGCTTCCCTTGC Rev: TCAGGATCTCCTCGTTCTGAC |
| GPX3 | Fwd: ATTTGGCTTGGTCATTCTGG Rev: CCACCTGGTCGAACATACTTG |
| GPX4 | Fwd: TCTGTGTAATGGGGACGATGC Rev: TCTCTATCACCTGGGGCTCCTC |
| DIO1 | Fwd: CAGGCCCCTGGTGTTGAAC Rev: ATCTACGAGTCTCTTGAACTGGT |
| DIO2 | Fwd: TCAGGTAACAATTATGCCTCGGA Rev: GCTGAACCAAAGTTGACCACC |
| DIO3 | Fwd: CTCGACTACGCACAAGGGAC Rev: GGTGGGCTTCCTCGATGTA |
| TR1 | Fwd: CCTATGTCGCCTTGGAATGTGC Rev: ATGGTCTCCTCGCTGTTTGTGG |
| TR2 | Fwd: GTTCCCCACATCTATGCCATTG Rev: GGTTGAGGATTTCCCAAAGAGC |
| TR3 | Fwd: CTTTGCAAGATGCCAAGAAA Rev: TCATGGCCTCCCAGTTGT |
| Scl-R(MRSR1) | Fwd: TTCGTCCCTAAAGGCAAGA Rev: CATTCGCAGTCCATGTCCTA |
| Sep15 | Fwd: GCTGTCAGGAAGAAGCACAA Rev: TTTTCATCCGCAGACTTCAA |
| Scl-H | Fwd: GGAAGAAAGCGTAAGGCGGG Rev: GGTTTGGACGGGTTCACTTGC |
| Scl-I-1 | Fwd: CTACTCCTGACATACTTCGACCC Rev: CCACGACAATCCAAACCCAG |
| Scl-K | Fwd: TCCACGAAGAATGGGTAGGA  Rev: GCTTCTCAGAGCAGACATTTACCT |
| Scl-M | Fwd: TGACAGTTGAATCGCCTAAAGGAG Rev: AACAGCACGAGTTCGGGGTC |
| Scl-N | Fwd: ACCTGGTCCCTGGTAAAGGAGC Rev: GGTGATGTCAAGGAAGTAGTTGGC |
| Scl-O | Fwd: GGCTGCCCATACCTGTGA Rev: CGTGCTGTTCGCTGTGTC |
| Scl-P | Fwd: CCTTGGTTTGCCTTACTCCTTCC Rev: TTTGTTGTGGTG·TTTGTGGTGG |
| Scl-S | Fwd: CAGAAGATTGAAATGTGGGACAGC Rev: CCTTTGGGGATGACAGATGAAGTAG |
| Scl-T | Fwd: TGAGGCTCCTGCTGCTTC Rev: GGTGGCGTACTGCATCTTTAAT |
| Scl-V | Fwd: TTGGGAATTGATGCTCCAGG  Rev: GCCTTCGGGTGAGTAGTTTC |
| Scl-W | Fwd: GGTGCCTCCCCAGAATCTAC Rev: TGGGGGAATTCAGAGAGAGA |
| SPS2 | Fwd: ACCGACTTCTTTTACCCCTTGG Rev: TCACCTTCTCTCGTTCCTTTTCAC |

**Supplemental Table 5. Left ventricular gene expression normalized to 36B4 (fold change)**

|  | **Se-0.15 ppm (control)** | | **Se-deficient** | |
| --- | --- | --- | --- | --- |
| **LV RNA expression levels** | **Males, n=5** | **Females, n=5** | **Males, n=5** | **Females, n=5** |
| ***Nppa*** | 1.03 ± 0.13 | 1.15 ± 0.28 | 1.77 ± 0.18 | 0.64 ± 0.12 |
| ***Nppb*** | 1.04 ± 0.13 | 1.02 ± 0.10 | 2.7 ± 0.26^****^ | 1.23 ± 0.18 |
| ***Gata4*** | 1.03 ± 0.12 | 1.01 ± 0.08 | 1.76 ± 0.20^**^ | 0.92 ± 0.06 |
| ***Gpx1*** | 1.07 ± 0.12 | 1.00 ± 0.02 | 0.63 ± 0.10 | 0.78 ± 0.06 |
| ***Gpx2*** | 1.01 ± 0.07 | 1.00 ± 0.03 | 0.72 ± 0.07^*^ | 0.71 ± 0.07^#^ |
| ***Gpx3*** | 1.14 ± 0.30 | 1.01 ± 0.07 | 1.33 ± 0.23 | 0.84 ± 0.13 |
| ***Gpx4*** | 1.02 ± 0.08 | 1.01 ± 0.06 | 1.04 ± 0.05 | 0.92 ± 0.07 |
| ***Dio2*** | 1.02 ± 0.11 | 1.02 ± 0.09 | 1.40 ± 0.29 | 0.80 ± 0.08 |
| ***Txnrd1*** | 1.02 ± 0.11 | 1.01 ± 0.06 | 1.09 ± 0.11 | 0.92 ± 0.10 |
| ***Txnrd2*** | 1.02 ± 0.09 | 1.01 ± 0.06 | 1.08 ± 0.07 | 0.92 ± 0.11 |
| ***Txnrd3*** | 1.02 ± 0.09 | 1.01 ± 0.06 | 1.10 ± 0.07 | 0.94 ± 0.12 |
| ***Msrb1*** | 1.01 ± 0.05 | 1.01 ± 0.07 | 0.95 ± 0.01 | 1.00 ± 0.05 |
| ***Selenof(Sep15)*** | 1.01 ± 0.07 | 1.00 ± 0.03 | 0.87 ± 0.04 | 0.70 ± 0.02^###^ |
| ***Selenoh*** | 1.09 ± 0.21 | 1.05 ± 0.15 | 0.41 ± 0.04^**^ | 0.60 ± 0.09 |
| ***Selenoi*** | 1.03 ± 0.12 | 1.03 ± 0.19 | 1.66 ± 0.16^*^ | 0.96 ± 0.07 |
| ***Selenok*** | 1.01 ± 0.06 | 1.00 ± 0.02 | 1.06 ± 0.02 | 1.00 ± 0.06 |
| ***Selenom*** | 1.02 ± 0.11 | 1.01 ± 0.08 | 0.62 ± 0.03^**^ | 0.82 ± 0.08 |
| ***Selenon*** | 1.31 ± 0.45 | 1.01 ± 0.07 | 1.83 ± 0.26 | 0.77 ± 0.05 |
| ***Selenoo*** | 1.15 ± 0.30 | 1.03 ± 0.15 | 1.61 ± 0.16 | 1.39 ± 0.21 |
| ***Selenop*** | 1.02 ± 0.09 | 1.00 ± 0.04 | 0.96 ± 0.07 | 0.72 ± 0.08^#^ |
| ***Selenot*** | 1.26 ± 0.44 | 1.01± 0.06 | 1.63 ± 0.34 | 1.06 ± 0.07 |
| ***Selenow*** | 1.01 ± 0.08 | 1.00 ± 0.05 | 0.17 ± 0.01^****^ | 0.35 ± 0.02^####^ |
| ***Selenos*** | 1.00 ± 0.03 | 1.00± 0.04 | 1.00 ± 0.07 | 0.81 ± 0.03^#^ |
| ***Sephs2*** | 1.14 ± 0.24 | 1.00 ± 0.04 | 1.34 ± 0.25 | 1.39 ± 0.04 |

**Supplemental Table 5. Left ventricular gene expression normalized to 36B4 (fold change).** Data are presented as fold change ± SEM relative to the corresponding sex-matched selenium-sufficient control group. For each group, n = 5 independent biological replicates, with each biological sample measured in technical duplicate by RT-qPCR. Statistical analyses were performed on ΔCt values using two-way ANOVA followed by Šídák’s multiple-comparisons test. Asterisks indicate comparisons between selenium-deficient and selenium-sufficient males, and hash symbols indicate comparisons between selenium-deficient and selenium-sufficient females. *P < 0.05; **P < 0.01; ****P < 0.0001; #P < 0.05; ##P < 0.01; ###P < 0.001; ####P < 0.0001.

**Supplemental table 6. RNA sequencing reveals expression patterns of cardiac hypertrophy in response to the Selenium-deficient diet in left ventricular tissue *(separate .xls file)***

RNA sequencing of isolated left ventricular tissue was performed. For the sequencing effort we used tissues of 10 animals which received control diet and 10 animals which received the selenium-deficient diet. In summary we found 2259 genes to be differentially expressed (DEGs; Padj <0.05) as the result of selenium deficiency. 1314 (58.2%) of these were found upregulated and 945 (41.8%) were downregulated with selenium deficiency. To reduce potential false positive signals, we applied a cut-off of logFC>|0.5|, reducing the DEGs to a total of 347 (145 upregulated and 202 downregulated)

**Supplemental table 7. List Antibodies.**

| **Antibody target** | **Company** | **Catalog number** | **Dilution** |
| --- | --- | --- | --- |
| GPX1 | Novus Biologicals | NBP1-33620 | 1:500 |
| GPX4 | Bio-techne | MAB5457 | 1:500 |
| Catalase | Protein tech | 21260-1-AP | 1:1000 |
| SOD2 | Protein tech | 24127-1AP | 1:3000 |
| SIRT1 | Cell Signaling | 8469 | 1:1000 |
| FOXO1 | Cell Signaling | 2880 | 1:1000 |
| Acetyl-FOXO1 | Invitrogen | PA5-104560 | 1:1000 |
| PER2 | Invitrogen | PA5-89045 | 1:1000 |
| **Secondary Antibody** | **Company** | **Catalog number** | **Dilution** |
| Goat anti-Rabbit | Dako/Agilent | P044801 | 1:2000 |
| Rabbit anti-mouse | Dako | P0260 | 1:2000 |
